# Crystallographic fragment screening and deep mutational scanning of Zika virus NS2B-NS3 protease enable development of resistance-resilient inhibitors

**DOI:** 10.1101/2024.04.29.591502

**Authors:** Xiaomin Ni, R. Blake Richardson, Andre Schutzer Godoy, Matteo P. Ferla, Caroline Kikawa, Jenke Scheen, William W Hannon, Eda Capkin, Noa Lahav, Blake H Balcomb, Peter G Marples, Michael Fairhead, Siyi Wang, Eleanor P Williams, Charlie W.E. Tomlinson, Jasmin Cara Aschenbrenner, Ryan M Lithgo, Max Winokan, Charline Giroud, Anu V Chandran, Martin Walsh, Warren Thompson, Jesse D Bloom, Haim Barr, Karla Kirkegaard, Lizbé Koekemoer, Daren Fearon, Matthew J. Evans, Frank von Delft

## Abstract

The Zika viral protease NS2B-NS3 is essential for the cleavage of viral polyprotein precursor into individual structural and non-structural (NS) proteins and is therefore an attractive drug target. Generation of a robust crystal system of co-expressed NS2B-NS3 protease has enabled us to perform a crystallographic fragment screening campaign with 1076 fragments. 47 fragments with diverse scaffolds were identified to bind in the active site of the protease, with another 6 fragments observed in a potential allosteric site. To identify binding sites that are intolerant to mutation and thus suppress the outgrowth of viruses resistant to inhibitors developed from bound fragments, we performed deep mutational scanning of NS2B-NS3 protease. Merging fragment hits yields an extensive set of ‘mergers’, defined as synthetically accessible compounds that recapitulate constellations of observed fragment-protein interactions. In addition, the highly sociable fragment hits enable rapid exploration of chemical space via algorithmic calculation and thus yield diverse possible starting points that maximally explore the binding opportunities to NS2B-NS3 protease, facilitating its resistance-resilient antiviral development.

## Introduction

Zika virus (ZIKV) belongs to the *Flaviviridae* family and is closely related to other flaviviruses such as Dengue virus (DENV), West Nile virus (WNV) and Yellow Fever virus (YEV). Although ZIKV infections typically manifest with mild symptoms, outbreaks in Brazil and French Polynesia revealed that the virus can cause microcephaly in new-borns and Guillain-Barre syndrome in adults^1–3^. While this development sparked significant global health concern, no vaccine or antiviral therapeutic is available for the prevention or treatment of ZIKV infection^4^.

The single-stranded, positive-sense ZIKV RNA is translated to produce a single polyprotein precursor that is cleaved by host proteases and viral protease NS2B-NS3 into individual functional proteins. NS3 contains a protease active site, with a serine, histidine and aspartate catalytic triad. Approximately 40 amino acids (aa) of NS2B function as a co-factor of NS3. These residues wrap around NS3, with the C-terminal residues forming a β-hairpin structurally contributing to the protease active site as well as participating in substrate recognition^5^. Structural studies have revealed several conformations of this two-component NS2B-NS3 protease, specifically a closed conformation^6^, an open conformation^7, 8^, and a super-open conformation^9^, depending on the dynamic interactions between NS2B and NS3. In the open and super-open conformations, the C-terminal portion of NS2B is disordered and distant from the NS3 catalytic site, resulting in an unstructured substrate binding site. These two conformations therefore have been considered as inactive forms, while the closed conformation of NS2B-NS3 with a well-structured binding pocket represents a promising targeting opportunity for antiviral development.

Two strategies have been deployed to target the NS2B-NS3 protease. The first involves the development of specific inhibitors that compete with the substrate on the active site. The second strategy focuses on non-competitive inhibitors, which target allosteric sites. Despite these efforts, neither competitive nor non-competitive inhibitors have advanced to clinical trials. Most of the reported competitive NS2B-NS3 inhibitors are limited to peptide mimics, such as macrocyclic inhibitors^10, 11^. Although these peptidomimetic inhibitors display nanomolar potency against ZIKV NS2B-NS3 protease, and some additionally have broad-spectrum antiviral activity, inhibiting growth of Dengue and West Nile viruses as well as Zika virus^12^, their cellular activity and pharmacokinetic properties are poor. In parallel, great effort has focused on developing allosteric inhibitors that block the interactions between NS2B and NS3 and lock the protease in the inactive form^13, 14^. This strategy has yielded several potent non-competitive inhibitors, including NSC135618^15^, which shows antiviral activity in cellular assays. Nevertheless, the allosteric site as well as the allosteric inhibitor binding mode have not yet been properly validated due to a lack of reliable structural information.

Viruses can evolve resistance to any inhibitor if the amino acids that such an inhibitor binds exhibit sufficient tolerance to mutation whilst the mutations that abrogate inhibitor binding do not greatly impair viral fitness. To prioritize NS2B-NS3-binding fragments that exhibit genetic constraint and thus are less likely to be able to evolve resistance, we performed deep mutational scan (DMS) of the NS2B-NS3 protease. DMS is a high-throughput experimental technique that relies on deep sequencing technology to measure the effect of each possible amino acid mutation at each position of a protein. DMS has emerged as a powerful tool in virology, particularly in studies of viral fitness and evolution^16^. Recent studies of the mutational potential of viral proteins, such as the SARS-CoV-2 spike protein^17^ and the ZIKV envelope protein^18^, provide rich information about the sequence-function relationships that apply to antibody engineering as well as vaccine design.

Here we combined crystallographic fragment screening with DMS to accelerate the development of resistance-resilient antiviral therapeutics targeting ZIKV NS2B-NS3 protease. Our crystallographic fragment screening against 1076 fragments identified diverse chemical scaffolds at both the active site and a potential allosteric site, providing structural details of fragment engagement for rapid follow-up compound design. The DMS results reveal the mutational potential of the hotspots for ligand engagement, which will guide rational compound design to suppress drug resistance at early stages of medicinal chemistry.

## Results

### Establishing a suitable crystal system for fragment screening

Several constructs have been reported to efficiently generate the ZIKV NS3 protease bound to the NS2B co-factor region. These approaches include the insertion of a glycine linker between the NS2B and NS3, as shown in the construct named gZiPro (Figure 1A), and the co-expression of the NS2B co-factor and NS3 protease in a bicistronic vector, as demonstrated in the construct named bZiPro^6^ (Figure 1A). Construct gZiPro has been commonly used for ZIKV NS2B-NS3 protease studies. However, the artificial glycine-rich linker raises concerns of altering the substrate-binding behaviour^19^. Therefore, the unlinked binary NS2B-NS3 protease is preferred for studies. In addition, bZiPro succeeded in producing a closed conformation of ZIKV NS2B-NS3 structure (PDB ID: 5GPI)^6^.

To produce crystals of ZIKV NS2B-NS3 protease in closed conformation for fragment screening, we generated a modified version of the PDB structure 5GPI^6^. The model of 5GPI contains 4 protein molecules in the asymmetric unit, with the N-terminal residues of NS3 in one molecule binding to a neighbouring active site, making this unsuitable for crystal soaking. A further issue with this system was the low expression yields of construct bZiPro in our hand. To address these issues, we generated a new construct, designated cZiPro (Figure 1A). In this construct, we removed the disordered C-terminal residues of NS2B and N-terminal residues of NS3 in 5GPI structure. This optimized construct yielded high quantities of ZIKV NS2B-NS3 protease and reproducibly produced crystals diffracting at 1.6 Å. Our NS2B-NS3 structure was determined in P4322 space group with only one protein molecule in the asymmetric unit, which is distinct from 5GPI model, yet the superposition of both structures revealed a backbone root-mean-square deviation (RMSD) of around 0.2Å (Figure 1B). The active site in our structure was not occluded by crystal packing, and is accessible via crystal soaking, thus making it suitable for fragment screening.

**Figure 1.**
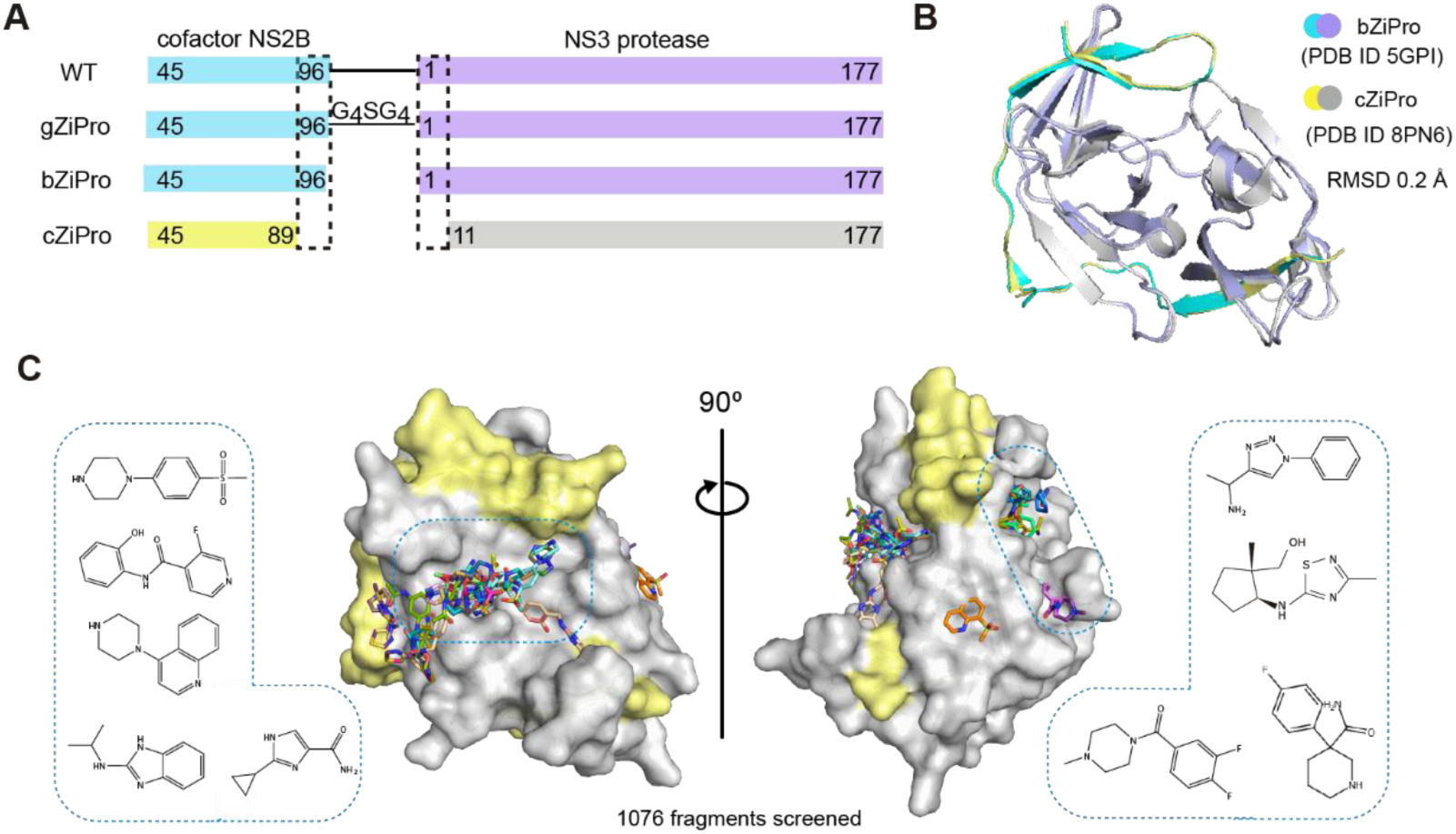
Crystallographic fragment screening of ZIKV NS2B-NS3 protease. (A) Domain boundaries of constructs gZiPro, bZiPro and cZiPro. (B) Structure superimposition of bZiPro structure 5GPI and cZiPro structure 8PN6 with a RMSD of 0.2 Å. (C) Surface view of ZIKV NS2B-NS3 fragment screening output in two orientations. NS3 protein surface is coloured in grey and NS2B is coloured in yellow. Identified fragments are shown as sticks. Examples of observed fragments are listed in dashed box.

### Fragment hit identification by crystallographic fragment screening

We used the newly established crystal system for a crystallographic fragment screening campaign against 1076 fragments using the XChem facility at Diamond Light Source. A total of 1058 datasets with resolution ranging from 1.4-2.3 Å were collected. 52 fragments were identified from these screening efforts (Crystallographic Supplementary Table), with 47 fragments bound in the active site and 6 in a potential allosteric site (one fragment bound to both the active site and allosteric site). Diverse chemical scaffolds were identified, including benzamide, benzothiazole/ benzimidazole and piperazine based fragments. Other moieties such as amides, pyrazoles, and sulfones were also observed from screened fragments (Figure 1C).

### Interaction motifs for ligand engagement in the active site

The active site of NS2B-NS3 protease site is divided into subsites S1, S1’, S2 and S3 based on previous study^6^ (Figure 2A). Of the 47 fragments observed in the active site, 44 bound in the S1, 1 in the S1’ and 2 in the S2, respectively, suggesting subsite S1 is a hotspot for fragment binding. In the S1 site, fragments, such as Z336080990(x0089), were frequently observed to form π-stacking with the side chain of Tyr161. The fragment was further stabilised by three hydrogen bonds, interacting with the side chain of Asp129, backbone of Tyr130, and a water molecule, respectively (Figure 2B). Fragment Z396117078(x1098) accommodated at both S1 and S1’ site (S1-S1’), forming two π-π stacking interactions with the side chains of Tyr161 and His51. In addition, its amide motif formed a hydrogen bond with the backbone of Gly151 as well as a water mediated hydrogen bond with the side chain of Tyr161 (Figure 2C).

S2 site binder piperazine fragment Z425338146(x0404) formed an electrostatic interaction with Asp83 from NS2B, while its benzene ring formed a π-stacking interaction with His51 (Figure 2D). We also observed one fragment Z1587220559(x0846b), which accommodated in S1’ site, forming two hydrogen bonds with backbones of Val36 and Val52 respectively (Figure 2E). Its benzimidazole ring formed hydrophobic interaction with surrounding residues, such as Val36, Leu30, and Met51 from NS2B.

Analysis of fragment-protein interactions revealed that π-stacking with the side chain of Tyr161 is a key interaction in the S1 site, represented by 39 of 47 fragment hits observed in this site (Figure 2F). Residues Asp129 and Tyr130 were also observed to form hydrogen bonds with nearly a quarter of identified fragments, together with Tyr161 and Tyr150 making the S1 site a hotspot for ligand engagement. In addition, residue His51, Ser135 and Asn152 from NS3, plus Asp83 from NS2B were also observed in fragment interactions (Figure 2G), indicating their potential contribution to ligand engagement.

**Figure 2.**
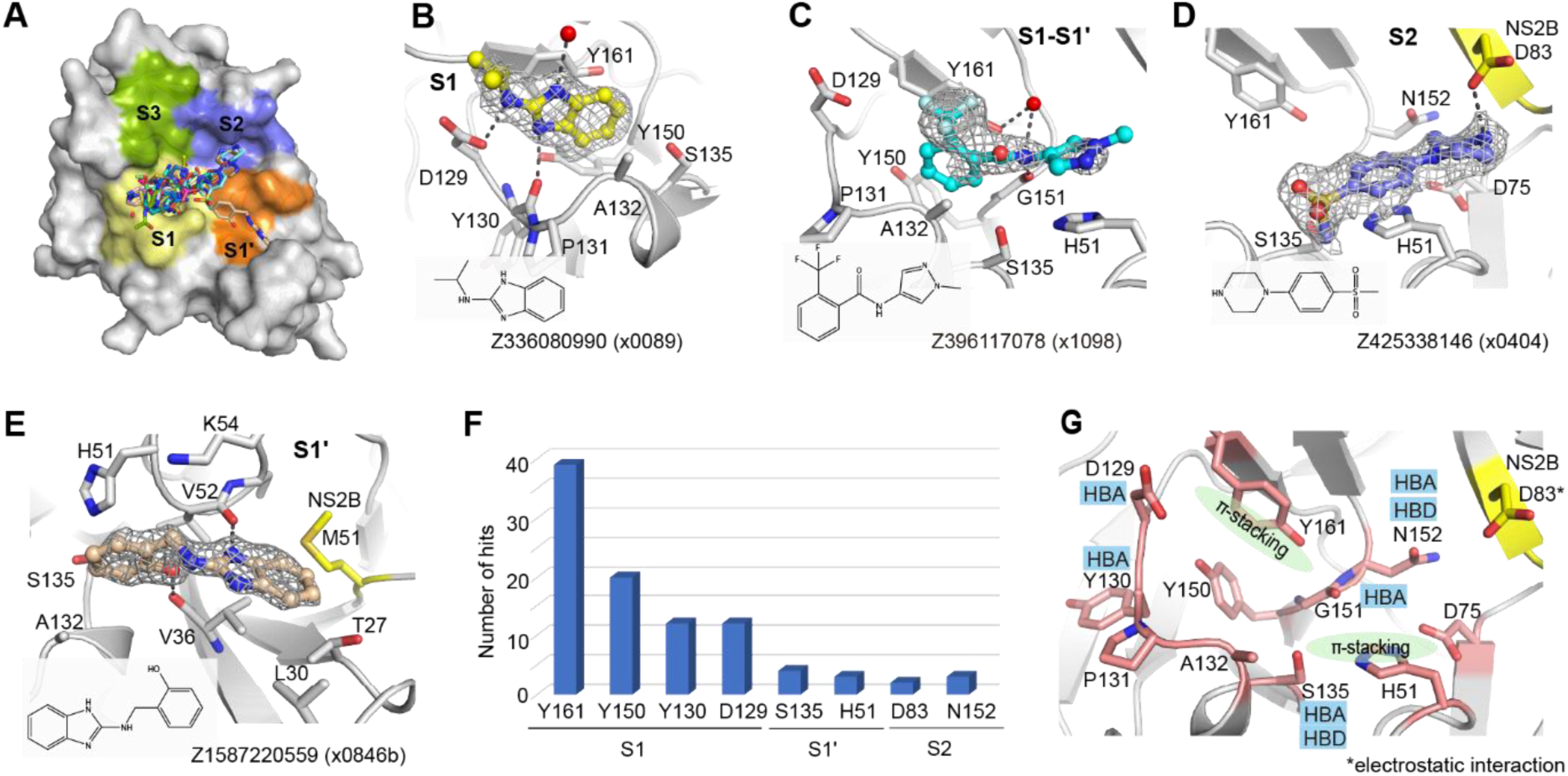
Interaction motifs for ligand engagement in the active site of ZIKV NS2B-NS3 protease. (A) Surface view of observed fragments bound in the active site. The active site is divided into subsites S1, S1’, S2 and S3 based on substrate residue binding position. Representative examples of fragments Z336080990(x0089) (B), Z396117078(x1098) (C) and Z425338146(x0404) (D) interacting at the S1, S1-S1’, and S2 site, respectively. The PanDDA event map is shown as a dark grey mesh. Hydrogen bonds are shown as dashed lines. (E)The unique fragment Z1587220559(x0846b) observed in S1’ site. (F) A plot of key residues observed for fragment interaction. Y-axis represents the number of fragments forming interaction with associated residues. (G) Key interacting residues revealed from fragment screening. HBA: hydrogen bond acceptor. HBD: hydrogen bond donor.

### Deep mutational scanning to measure mutational tolerance of ZIKV NS2B-NS3 protease

Given that viral proteases often evolve resistance to inhibitors through mutations in key residues, understanding the mutational tolerance of ZIKV NS2B-NS3 protease is essential for designing resistance-resilient antivirals. To gain a comprehensive view of how specific amino acid substitutions, especially at key ligand interacting residues, affect the protease’s function and ligand engagement, we employed deep mutational scanning (DMS). Specifically, we generated libraries encoding all single amino acid substitution mutants of the entire coding sequence of ZIKV NS2B-NS3 using a previously reported infectious clone of the ZIKV prototype strain MR766^20^. To ensure comprehensive coverage of all possible mutations during library cloning and screening stages, we divided the NS2B-NS3 coding sequence into three distinct ‘tiles’ (Supplementary Figure 1A) and created multiple mutant plasmid libraries for each tile (three each for tiles 1 and 2, and two for tile 3), which were handled separately in all subsequent steps to provide true biological replicates. Each library contained an average of 2.7x10^6^ unique plasmid clones, which overrepresents the total number of possible codon variants of 3,296 (32 combinations of NNK multiplied by 103 codons) by 819-fold, with an average of 1.05 mutations per clone.

Mutant virus pools were produced by transfection of HEK 293T cells with the plasmid DNA libraries and selected for viral fitness by passaging through Huh-7.5 cells^21^ (Supplementary Figure 1B). We used deep sequencing to quantify the frequency of each mutation in the mutant viruses relative to the initial plasmid mutant libraries^22^. Stop codons, which are expected to be uniformly deleterious, were greatly purged, and nonsynonymous mutations, many of which will be deleterious, decreased in frequency (Supplementary Figure 1C).

We next estimated the preference of each site in the NS2B-NS3 protease protein for each amino acid. These preferences represent enrichments of each amino acid at a site after selection for viral growth, normalized to the abundance of the wild type codon at each site. Although there is some noise, the preferences were strongly correlated among three replicates (Supplementary Figure 1D). We experimentally quantified the relative fitness of a panel of individual NS2B-NS3 mutants and found these values highly correlated (Pearson correlation of 0.769) with the mutation effects measured by DMS (Supplementary Figure 1E).

The mutational tolerance at any given codon can be determined from the effects of each individual mutation at that site on viral fitness. The across-replicate average of the effects of each mutation at each site in the NS2B-NS3 protease protein was calculated as the log-fold change in the frequency normalized to the change in frequency of a wild type control and represented in the heatmap in Figure 3A. At the lower end of fitness, stop-codon mutations and catalytic triad mutations are considered to be deleterious. Thus, their mutational effects have been applied as reference sets within the distribution of fitness effects of NS2B-NS3 mutations (Figure 3B). At the upper end of the fitness distribution, we set a threshold of fitness at -1.0 (Figure 3B). Mutant viruses that have a mutational effect of more than -1.0 are tolerated and allow the growth of virus, even though the fitness is reduced from that of wild-type. A mutational effect of more than zero could be considered hyper functional, suggesting that such mutation is beneficial for virus fitness under the growth conditions of the assay.

Most mutations in NS2B-NS3 reduce the fitness of the virus in Huh-7.5 cells (Figure 3A, blue squares), suggesting low mutational tolerance at many residues. Many residues, however, such as NS2B residues 88-94 and NS3 residues 12-20, exhibit high tolerance for mutation (white squares). Notably, mutations at these sites do not decrease viral fitness in Huh-7.5 cells, instead displaying fitness similar to that of viruses with wild type residues (X) at these positions. Some sites, such as NS3 Arg106 (pink squares), showed both a high tolerance for changes and an increase in fitness when certain mutations were introduced. Residues in the protein that have a high tolerance for mutations are not good candidates for ligand binding sites, because mutations that disrupt ligand binding do not compromise viral fitness.

To facilitate analysis of these data, we generated a visualization scheme, termed ‘Fitness View’, to map the mutational tolerance of each residue to our crystal structure (Figure 3C). Surfaces that are coloured in white indicate sites that are mutationally intolerant, while residues that are dark red represent a high tolerance for mutations. Overall, this fitness map provides a wealth of information about the sequence-function relationships in the NS2B-NS3 protease protein that can be used to guide the selection of key interacting residues for rational antiviral design.

**Figure 3.**
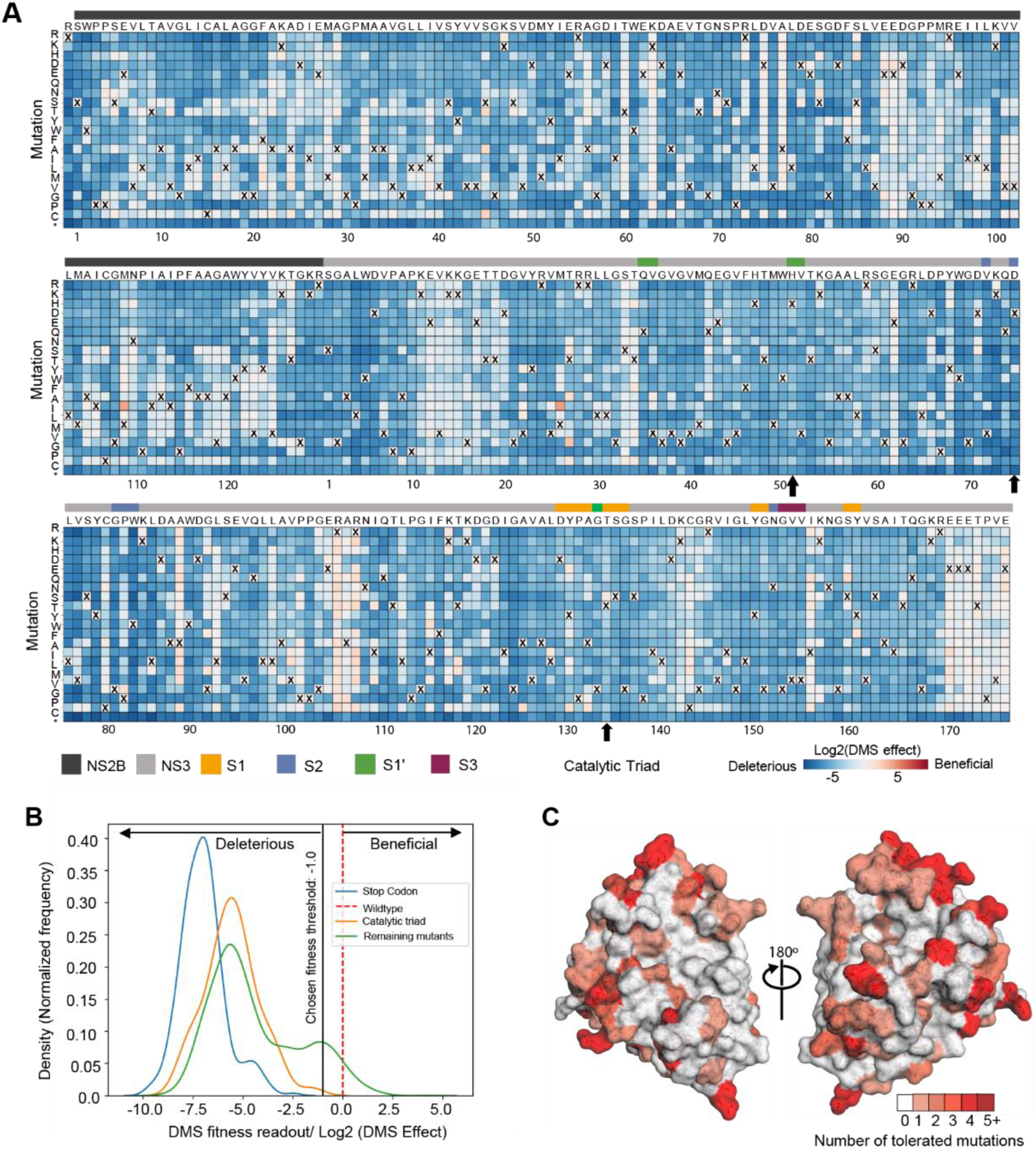
Deep mutational scanning to measure mutational tolerance of ZIKV NS2B-NS3. (A) The heatmap of NS2B-NS3 residues indicating the mutational effect of each amino acid substitutions in the ZIKV NS2B-NS3 protease. Blue mutations are deleterious for viral growth in Huh-7.5 cells relative to wildtype, white mutations are neutral, and red mutations increase growth in Huh-7.5 cells. Wildtype amino-acid identities at each site are denoted by an ‘X’. In general, most mutations decrease the fitness of the virus in Huh-7.5 cells. (B) Distribution of fitness estimates for all mutations. The distributions of fitness effects of stop-codon mutants, Catalytic triad mutants and the remaining mutants are coloured in blue, orange and green, respectively. The black vertical line indicates the threshold -1.0 (Log2 (DMS effect)) for fitness. (C) Fitness view of NS2B-NS3 protease. Mutational tolerance is mapped onto the structure of ZIKV NS2B-NS3 (PDB ID: 8PN6). Residues marked in white do not tolerate changes to that site, while residues marked in red tolerate a range of changes. The number of changes tolerated is indicated by the number and the colour.

### Fitness view reveals potential mutational effect on ligand engagement in the active site

Inspection of the fitness view in the active site reveals that S1’ and S2 subsites are relatively intolerant of nonsynonymous mutations (Figure 4A), while a few positions in the S1 subsite and at the C-terminus of NS2B show mutational tolerance, indicating a risk of potential resistance mutations (Figure 4A). Specifically, Tyr130 in S1 subsite can potentially mutate to multiple amino acids, such as cysteine, glutamic acid. At position 132, cysteine and proline substitutions are predicted to have similar replication efficiency to the wild type alanine (Figure 4B). Despite the high mutability of Tyr130 and Ala132, S1 fragment binders from our screen predominately interact with the backbone of Tyr130 and indirect contact with Ala132. Thus, the effect of the potential mutations of Tyr130 and Ala 132 on ligand engagement is estimated to be limited. Similarly, mutational flexibility of Gly82 and Phe84 at the C-terminus of NS2B is less concerning. Phe84 hasn’t been observed for fragment interaction, while Gly82 utilised its backbone for fragment interaction.

However, the potential mutation of tyrosine to phenylalanine at position 161 of NS3 raises our concern. Although Tyr161 contributes to fragment interaction mainly via π-stacking, which can be replaced by phenylalanine, we also observed one fragment Z120319681(x0719) that formed a direct H-bond with the hydroxyl group of Tyr161. Mutating tyrosine to phenylalanine would abolish the hydrogen bond formation. Thus, hydrogen bonding to Tyr161 is suggested to be avoided for follow-up compound design to decrease the probability of drug resistance. In contrast, key interacting residues Asp129 in the S1 subsite and Asp83 (NS2B) in S2 subsite are mutational intolerant, suggesting opportunities for fragment growth (Figure 4A, 4E).

**Figure 4.**
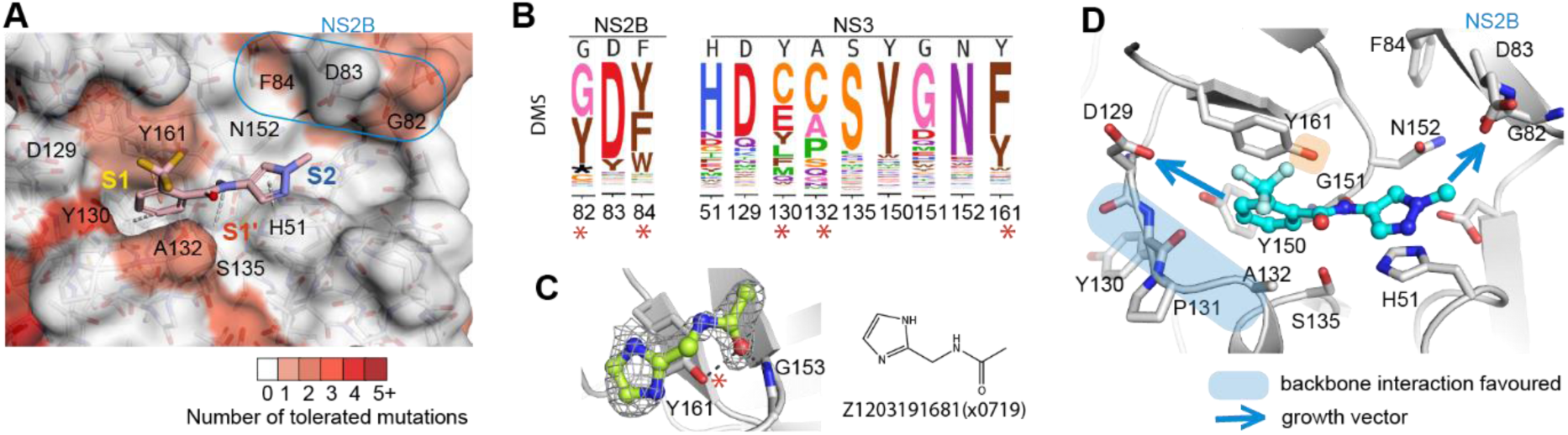
Fitness view reveals potential mutational effect on ligand engagement in the active site. (A) Fitness view of the active site with fragment Z396117078(x1098) presented. Key interacting residues identified from fragment screening are numbered. Mutational intolerant region is coloured in white, and mutational tolerant region is coloured in red. (B) Logo plot of experimentally measured amino acid preferences of the label key interacting residues in figure A. High mutational residues are marked with an asterisk sign. (C) Example of Fragment Z1203191681(x0719) formed a direct H-bond with the hydroxyl group of tyrosine 161. (D) Key interacting residues in the active site. Regions that suggest to form backbone interacting are coloured in blue. Side chain of Y161 is highlighted by colour orange due to its potential mutation to phenylalanine. Opportunity for fragment growth is shown as an arrow.

### Fitness view invalidates the reported allosteric site as resistance-robust opportunity

Six fragment hits were observed outside of the orthosteric site, in a pocket located beneath the catalytic residues and next to the interface between the C-terminal β-hairpin of NS2B and NS3 (Figure 5A). This potential allosteric site contains two sub-pockets, named as allosteric site AS1’ and AS2’ (Figure 5B, 5C). AS1’ utilises Trp69 and Trp83 as its major interacting residues. One example, Z57122377(x0130), formed π-stacking with the side chain of Trp69 and was further stabilised by hydrogen bonding to the backbone of Trp83 (Figure 5B).

AS2’ is formed mainly by hydrophobic residues, including Phe116, Ile123, Ala164, and Ile165 from NS3, and Leu78 and Phe84 from NS2B. Four fragments with different scaffolds were observed in AS2’ with different poses. Fragment Z1428159350(x0806) contains a benzene ring that formed hydrophobic interactions with surrounding hydrophobic residues (Figure 5C), while the nitrogen atom N2 from its triazole ring formed a hydrogen bond with the backbone of Gln167. Another hydrogen bond was formed between its amine group and a water molecule (Figure 5C). In contrast, Fragment Z1272515105(x0777) bound in AS2’ exhibited a completely different binding mode. Its hydroxyl group formed two water-mediated hydrogen bonds with the side chains of Thr118 and Asp120 (Figure 5D). The N2 atom from its thiadiazole ring formed another water-mediated hydrogen bond with Asp71, and the other nitrogen atom (N5) formed a hydrogen bond with water from solvent. Its 5-carbon ring fit into a hydrophobic cavity that is formed by Ile123, Phe116 and Ala164. In conjunction, the fragments bound in this non-active site mapped a potential allosteric pocket next to the interface of the C-terminal β-hairpin of NS2B and NS3. This observation agrees with previous work of characterizing allosteric pockets in ZIKV NS2B-NS3 via docking and mutational studies^23, 24^ (Figure 5E).

We next investigated the genetic flexibility of the key interacting residues in the allosteric site. In comparison to the active site, the allosteric site has relatively higher mutability (Figure 5F). In particular, the mutational flexibility of residues Asp71 and Thr118 raise our concern. Asp71 was observed to interact with fragments via hydrogen bonding (Figure 5D). Its potential of mutating to serine would alter its charge as well as the length of the side chain, raising risk of maintaining interactions with ligands. Similarly, polar residue Thr118 has the potential of mutating to methionine, which has a longer side chain that potentially narrows the pocket, likely resulting in crashing with the ligand. Considering the potential mutational risk as well as underdefined key interacting residues in the allosteric site, we, therefore, prioritized our work on the active site of the protease.

**Figure 5.**
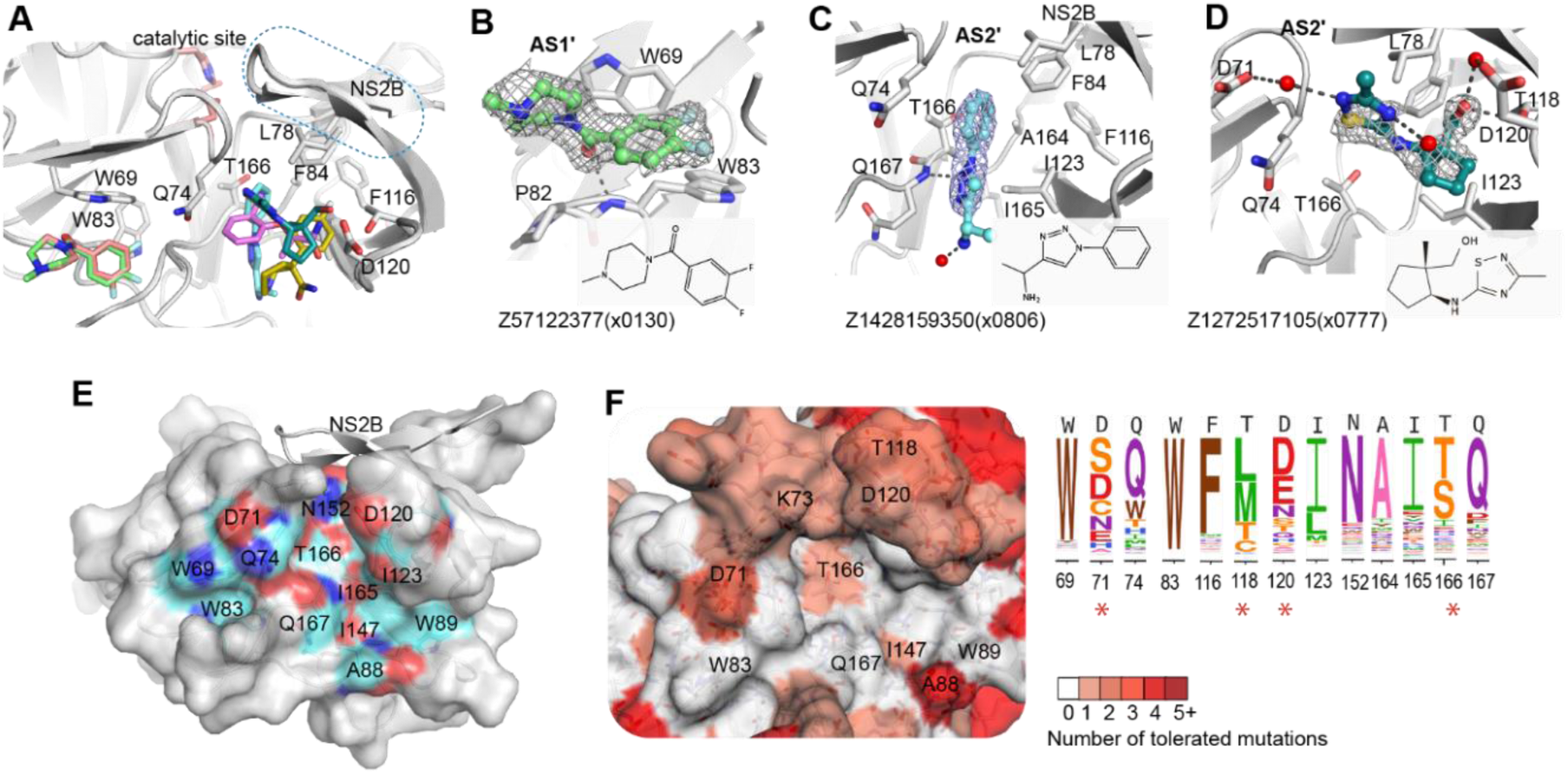
Fragments bound in a non-active site of ZIKV NS2B-NS3 protease. (A) Overview of fragments observed in the non-active site. (B) Structural details of observed fragment Z57122377(x0130) interacting at AS1’. Residues that participate in fragment interaction are shown as sticks. Representative examples of fragment Z1428159350 (x0806) (C) and Z1272517105 (x0777) (D) bound in the AS2’ site. Hydrogen bonds are shown as dashed lines. The PanDDA event map is shown as a dark grey mesh. (E) Surface view of an allosteric site defined by reported docking and mutational studies. Labelled residues are reported for ligand interaction. Oxygen, nitrogen and carbon atoms are coloured in red, blue and cyan, respectively. (F) Fitness view of the potential allosteric site mapping to Figure E. The logo plot presents the experimentally measured amino acid preferences of the interacting residues in the non-active site. High mutational residues are marked with an asterisk sign.

### Identified fragments are undetected in binding and activity assays

To further test whether our screened fragment binders have promising binding affinity and inhibitory activity, we set up a GCI-based CreoptixWave system for binding assay and a fluorescence-dose-response inhibitory assay^25^. A reported peptide-hybrid inhibitor compound **36**^26^ was used as the positive control (Figure 6D, 6H). However, we either failed to get conclusive results or measurable signal to confidently validate these screened fragments. Some fragments vaguely showed close to mM range affinity, such as fragments Z270834034 (x0472) and Z1269184613(x0772) (Figure 6A, 6B), but we failed to get consistent results during replicates. Most of the tested fragments had a nondetectable response signal or no binding activity, which showed a response signal near or below zero after the reference subtraction, such as fragment Z1587220559(x0846) (Figure 6C). Unsurprisingly, none of the screened fragments showed inhibitory activity in the biochemical assay (Figure 6E-G). Such observations are in common with many crystallographic fragment screens that only a low percentage of crystallographic fragment hits can be reliably detected in biophysical or biochemical assays^27^. This is also supported by a comparison study of different fragment screening assays, such as NMR, MST and enzyme inhibitory assay, showing limited overlapping of fragment hits^28^.

**Figure 6.**
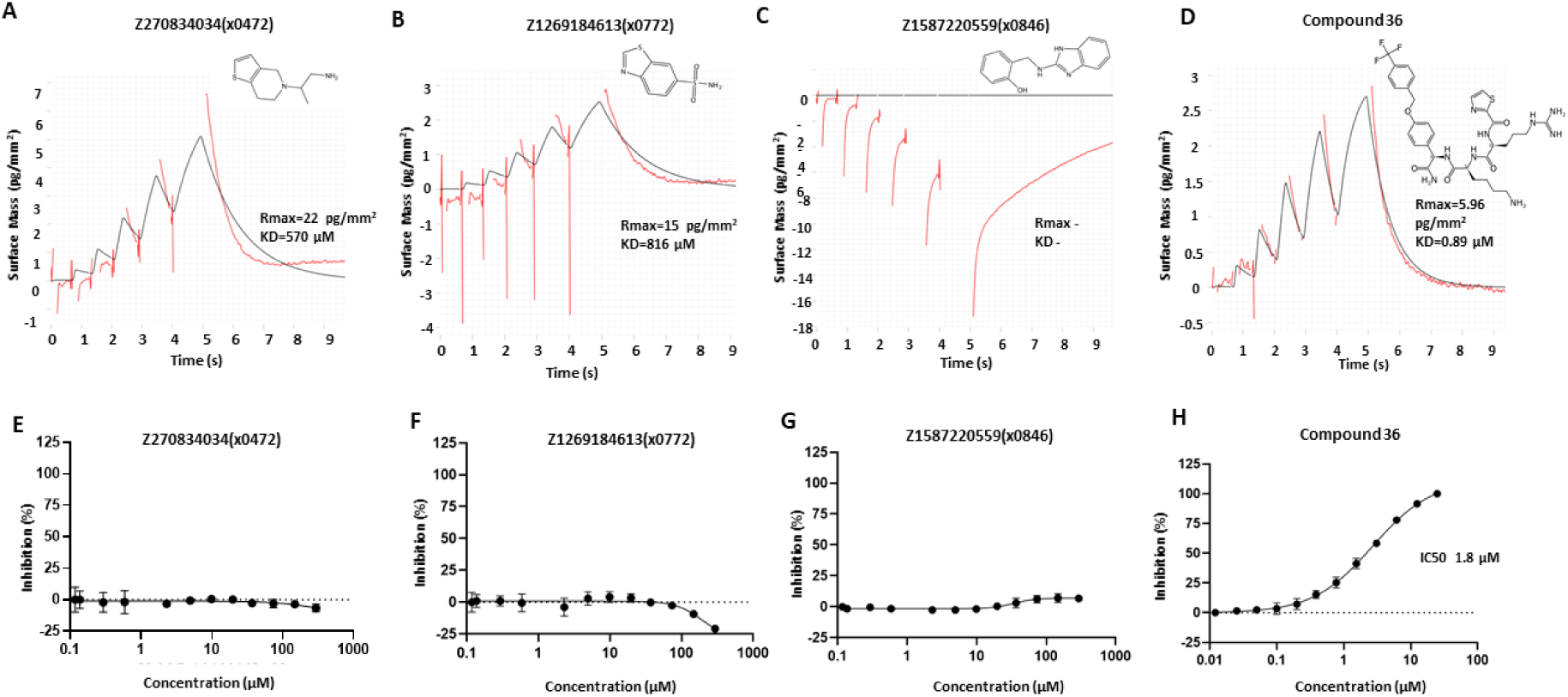
Screened crystallographic fragments tested in CreoptixWAVE binding assay and inhibitory activity assay. Fragment Z270834034(x0472) (A) and Fragment Z1269184613(x0772) (B) showed close to mM affinity, but such results could not be reproduced. (C) Fragment Z1587220559(x0846) selected as an example of negative result. (D) Peptide-hybrid inhibitor compound **36** was used as the positive control, showing a Kd of 0.89 µM in the GCI assay. Compound **36** was tested at 1 µM, while fragments were tested at 250 µM. 2D structures of compounds are shown on top. (E-H) Fluorescence-dose-response inhibitory activity assay. Fragments Z270834034(x0472) (E), Z1269184613(x0772) (F) and Z1587220559(x0772) (G) showed no inhibitory activity. (H) Positive control compound **36** showing an IC50 of 1.8 µM.

### Opportunities for rapid follow-up compound design via fragment merging and linking

With careful inspection of fragment-protein interaction profiling and fitness view in the active site, fragments Z1619958679(x0852), Z396117078(x1098), Z425338146(x0404), and Z1587220559(x0846b) were selected for merging (Figure 7A). Z1619958679(x0852) recaptured core interactions observed in the S1 site, specifically aromatic interaction with Tyr161, hydrogen bonding with the backbone of Tyr130. Z396117078(x1098) is the only fragment observed that bridges the S1 and S2 subsites. Its benzene ring superimposed with the benzene ring of fragment Z1619958679 (x0852) in the S1 subsite (Figure 7B, left), suggesting that these two fragments could be merged into a single, larger scaffold. In addition, Z396117078(x1098) partially overlapped with fragment Z425338146(x0404) in the S2 subsite (Figure 7B, middle). Merging Z396117078(x1098) and Z425338146(x0404) derived a pyrazole piperidine scaffold, which expanded the small molecule to engage the S1, S1’ and S2 simultaneously, represented by Enamine compound Z2451096209 (Figure 7C, middle).

In addition, the diverse fragments we observed in S1 site provide rich opportunities to grow pyrazole piperidine scaffold, which can potentially target key interacting residues such as the side chain of Asp129 and backbone of Tyr130. Enamine compound PV-004740099668 was presented as an example (Figure 7C, left). Such a pyrazole piperidine scaffold and S1 binder mergers were further assessed by algorithmic calculation called Fragmenstein^29^, which is based on the principle that the pose of mergers should preserve the poses of its parent fragments. We limited the conformational derivation of predicted mergers to be less than 1 Å in comparison with their parent hits in order to potentially maintain its parent fragments interactions. Such as the merger PV-004740099668 has 0.82 Å RMSD from its parent fragments Z1619958679(x0852) and Z396117078(x1098), and 0.27 Å for merger Z2451096209 respectively (Figure 7C).

**Figure 7.**
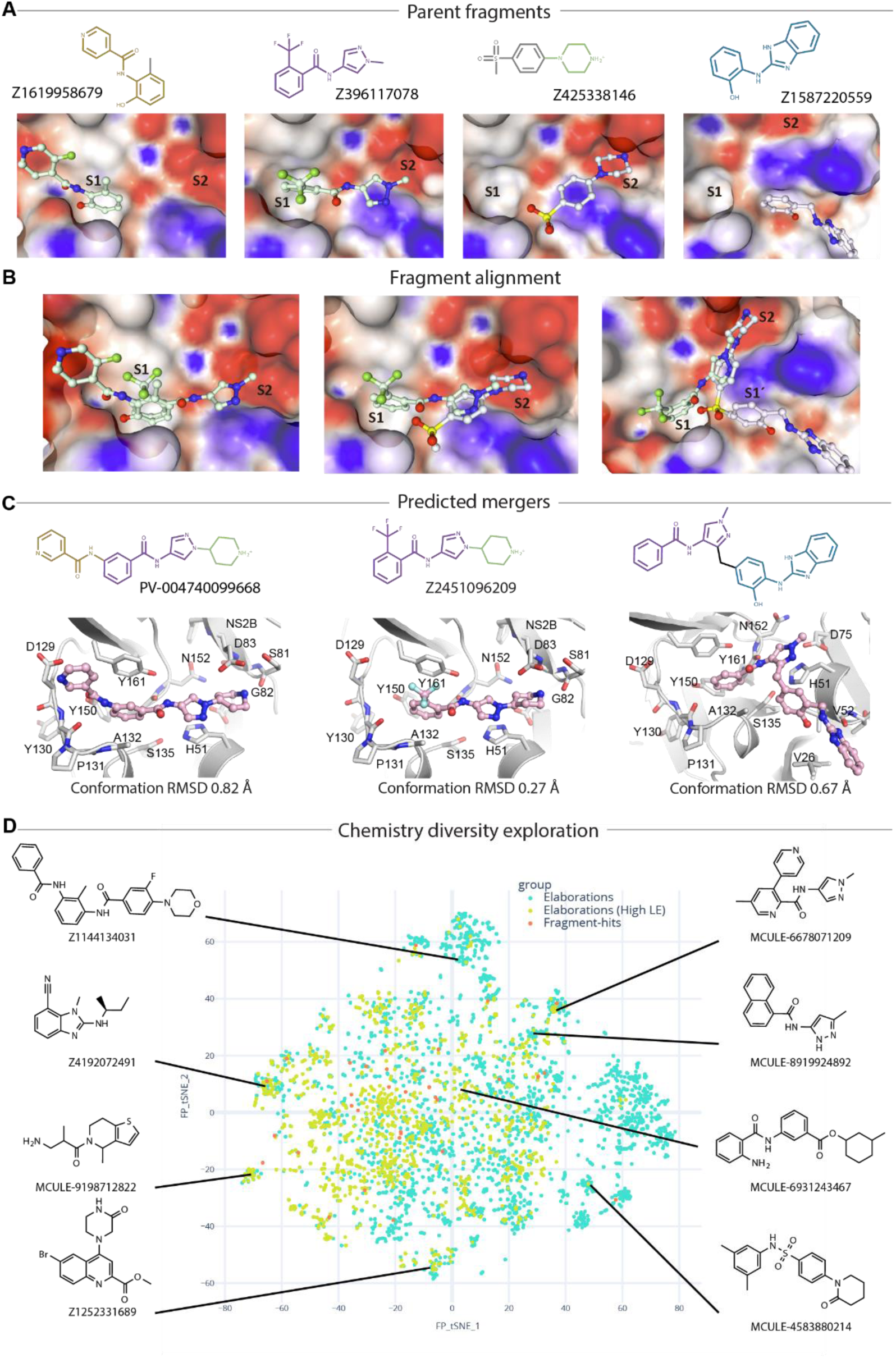
Fragment bridging multiple sites providing opportunities for merging and linking. (A) Fragments selected as parent hits for rapid follow-up compound design. (B) Fragment alignment shows opportunities for merging and linking. (C) Predicted mergers posed in the active site. 2D structure is shown at top with its Enamine compound identity code. RMSD value indicates the conformational variance to its parent fragments. (D) An annotated plot of the chemical diversity of 4,000 filtered catalogue compounds that are close analogues (graph edit distance fewer than 6 edits) of mergers of the active site fragment-hits searched with SmallWorld in Enamine and Mcule catalogues.

In addition to merging, we also observed that fragment Z1587220559(x0846b), which uniquely accommodated in the S1’ site, provides linking opportunities with fragments Z396117078(x1098) via a methyl linker without causing strain or internal clash (Figure 7C, right). This is supported by the RMSD value of 0.67 Å compared to its parent fragments while keeping a reasonable angle (112°) of the methyl linker. Such binding region has not been explored for small molecule engagement. Our observation, despite only one fragment, brings new opportunities for novel scaffold inhibitor development.

To further explore chemistry diversity, we used all screened fragments as input for merging via Fragmenstein. A pipeline of automated merging, catalogue searching and constrained docking revealed a large amount (around 4000 filtered catalogue compounds) and diversity of possible mergers that could be explored (Figure 7D), such as benzimidazole-based compound Z4192072491 (Enamine) and pyrazole naphthalene MCULE-8919924892. Both the mergers and the analogue docking were computed with Fragmenstein and the final compounds were predicted to be faithful to the parent fragments (RMSD < 1 Å) without violating the energy scoring function (ΔGcalc of binding < –5 kcal/mol). The abundant and diverse chemical matter suggested from algorithmically merging demonstrates that our screened fragments are highly sociable, enabling many possible avenues of exploration.

## Discussion

We obtained a robust ZIKV NS2B-NS3 crystal system that enabled a successful crystallographic fragment screen. The sampled fragment hits provide diverse possible starting points for rapid follow-up compound design. This brings new opportunities to this antiviral target where currently the orthosteric inhibitors are largely limited to peptidomimetics^26^. In addition, analysis of fragment-protein interaction profiling highlights the S1 subsite as the major targeting opportunity for orthosteric inhibitor development. This is supported by over 80% of observed fragments being accommodated in this region. Such a high rate of hits observed in S1 site is likely due to the deep pocket mainly formed by the bulky side chains of Tyr150 and Tyr161.

Drug resistance is one of the major challenges that an effective protease inhibitor needs to confront. Lessons have been well learnt from other drug discovery campaigns, such as drug resistance in HIV-1 protease^30^. Knowledge of the potential of the key interacting residues to tolerate mutations is necessary to design effective antivirals to decrease the probability of drug resistance. In this study, DMS has been applied to profile the mutational potential of our target protein. This helps to assess fragment-protein interactions, constraining the binding region for ligand engagement as well as highlighting the opportunities for fragment growth. We believe that incorporation of DMS at the very early stage of medicinal discovery is necessary and beneficial for effective compound design.

Multiple allosteric sites of Flaviviral NS2B-NS3 protease have been proposed but without structural description^9, 15, 31^. Our fragment screening data presented herein, helps to reduce this knowledge gap and map the allosteric site with structural details. Although the allosteric binding pocket we described here presents in a closed conformation, it agrees with a proposed allosteric site reported previously^9, 15^, supporting the ligandability of the interface of the C-terminal β-hairpin of NS2B and NS3. However, the distinct binding poses and interactions observed from our screening indicate the challenges for non-competitive inhibitor development. More importantly, DMS data revealed a high mutability of several interacting residues in this allosteric site, suggesting potential risks of drug resistance. We, therefore, prioritized the orthosteric site as the antiviral target. We hope this observation will provide novel insights into this field given that there is increasing effort in allosteric inhibitor development for Flavivirus NS2B-NS3 protease^26^.

Merging fragments can effectively lead to on-scale affinity from non-measurable weak binders^32^, and is therefore commonly applied for rapid follow-up compound design. This has been proven by several successful fragment-to-lead case studies, with targets including the SARS-CoV-2 main protease^25^ and NSP3 macrodomain^33^. In this study, all the screened crystallographic fragments failed to show confident binding affinity and inhibitory activity. This is not surprising as crystallographic fragment screen is a highly sensitive assay, likely due to the high concentration (100 mM in crystal drops) used in soaking. However, the structural details of fragment-protein interactions provide clear routes, or rich information to grow fragments for higher affinity. In our case, a key fragment Z396117078(x1098) bridges the S1 and S2 subsites and provides an essential pose for merging. Merging this fragment with S1 binders derives an easily accessible benzamide-(pyrazolyl) piperidine scaffold, supported by over 1000 analogues available from Enamine REAL. The easily accessible predicted mergers suggested here offer a potential route for rapid follow-up compound design.

Overall, this study has combined crystallographic fragment screening and deep mutational scanning of Zika virus NS2B-NS3 protease to accelerate the development of resistance-resilient inhibitors. Crystallographic fragment screening samples fragment hits and maximally explores the binding opportunities to NS2B-NS3 protease for fragment-to-lead development, and DMS results assist in prioritizing fragment-protein interactions for rational compound design.

## Methods

### Protein expression, purification and crystallization

A bicistronic construct here named cZiPro was based on the sequence of PDB model (PDB ID:5GPI). The construct was created by using a synthetic *E. coli* codon-optimised gene ZIKV sequence as template (accession YP 009227202.1). Golden gate cloning was used to insert this into the pNIC-HIS6-GST-TEV-GG vector^34^. The final construct contains a GST fusion with a TEV site followed by the NS2B peptide (residues 45-89 aa). This is followed by an intervening ribosome binding site and the NS3 protease domain (residues 11-177aa). Cells cultured in TB media were initially grown at 37 °C, and they were induced with 0.5 mM IPTG at 18°C overnight. Harvested cells were lysed by sonication in a buffer containing 50 mM HEPES, pH 7.5, 500 mM NaCl, 20 mM imidazole, 5% glycerol and 1 mM tris (2-carboxyethyl) phosphine (TCEP). The recombinant protein was initially purified by Ni^2+^-affinity chromatography. The His6 and GST tag were removed by TEV protease treatment, and the cleaved protein was passed through Ni^2+^ affinity beads and further purified by size exclusion chromatography using HPLC column in the buffer containing 25 mM HEPES pH 7.5, 150 mM NaCl, 0.5mM TCEP and 5% Glycerol. The co-expressed protein NS2B-NS3 was concentrated to ∼15 mg/ml for crystallization. Crystallization was performed using the sitting-drop vapor diffusion method at 20 °C. The crystals were obtained in the condition containing 30% w/v PEG2000, 0.2 M ammonium sulfate, 0.1 M sodium acetate, pH 4.8

### Crystallographic fragment screening, data collection and analysis

The fragment screening was performed using the XChem facility at Diamond Light Source, UK. The fragments from the DSi-Poised Library^35^, MiniFrags Probing Library^36^, CovHetFrags^37^, and SpotXplorer^38^ were dispensed into crystal drops by ECHO Liquid Handler with 20% (V/V) DMSO in the final condition and incubated for three hours at 20 °C. Crystals were harvested using Crystal Shifter ^39^ (Oxford Lab Technologies) and cryo-cooled in liquid nitrogen.

Diffraction data were collected at the I04-1 beamline at Diamond Light Source at 100 K and processed with automated pipelines using combined software, including XDS^40^, Autoproc^41^, Xia2^42^ and DIALS^43^. All further analysis was performed using XChemExplorer^44^. Ligand restraints were generated with ACEDRG^45^ and GRADE^46^. Fragment hit identification was analysed by PanDDA^47^. Electron density map was generated by Dimple^48^. Model building and refinement were carried out in COOT^49^ and Buster^50^ via XChemExplorer platform.

### Cell lines and antibodies

HEK 293T and Huh-7.5^21^ (provided by Charles M. Rice, Rockefeller University) cells were grown in Dulbecco’s modified Eagle’s medium (DMEM; Gibco BRL Life Technologies, Gaithersburg, MD) with 10% fetal bovine serum (FBS; Gibco BRL Life Technologies). The humanized monoclonal antibody D1-4G2-4-15 (4G2) (Absolute Antibody, Oxford, UK) is a broadly reactive flavivirus antibody that binds to an epitope at the fusion loop domain of the E protein^51^.

### Deep mutational scanning of ZIKV NS2B-NS3 protease

Here we present a compressed summary of the methods used for ZIKV NS2B/NS3 protease DMS and the validations of the results. Detailed methods can be found in Supplementary information_DMS experiment.

To generate mutant plasmid libraries, we created NS2B-NS3 protease codon-mutant DNA fragments using a previously described PCR mutagenesis approach^52^ with sets of forward and reverse oligos that randomized each codon with an NNK sequence where N is any nucleotide and K is either a G or a T. These products were cloned into our previously described single-plasmid reverse genetics system for ZIKV strain MR766 (sequence is available at Genbank accession KX830961)^20^ using techniques like those described in our previous ZIKV envelope protein DMS paper^18^. We generated infectious virus stocks of these ZIKV NS2B-NS3 protease DMS libraries by transfecting plasmids into HEK 293T cells using a protocol to maintain library complexity as previously reported^18, 20, 53^. Wild type virus was rescued in parallel as a control. These stocks were titered on Huh-7.5 cells, and then selected by infecting these cells at a MOI of 0.05 infectious units per cell. Infected cells were collected at day 2 post infection.

Total RNA was extracted from these cells, reverse transcribed, and subjected to Illumina deep sequencing using a barcoded-subamplicon sequencing approach described in previous work^54^ (see also https://jbloomlab.github.io/dms_tools2/bcsubamp.html) to minimize sequencing errors. The raw deep sequencing data have been deposited in the Sequence Read Archive as BioProject PRJNA1125458. The code that performs the analyses of the deep sequencing data is available on GitHub at https://github.com/jbloomlab/ZIKV_DMS_NS3_EvansLab. This repository includes summary notebooks that provide detailed statistics like read depth and mutation frequencies for each library tile (see subdirectories of https://github.com/jbloomlab/ZIKV_DMS_NS3_EvansLab/results/summary/). Briefly, we used dms_tools2^55^ (https://jbloomlab.github.io/dms_tools2/), version 2.4.14, to count the occurrences of each mutation in each sample (see https://jbloomlab.github.io/dms_tools2/bcsubamp.html for details). The amino-acid preferences were computed from these counts using the approach previously described^55^ (see also https://jbloomlab.github.io/dms_tools2/prefs.html). The mutational effects are the log of the preference for the mutant amino acid divided by the preference for the wild type amino acid. Output containing the numerical values of the counts of mutations in each sample, the amino-acid preferences, and mutational effects are processed on a per-tile basis. These data are provided in CSV file format in the GitHub repository. See the README (https://github.com/jbloomlab/ZIKV_DMS_NS3_EvansLab/blob/main/README.md) for details on navigating the analysis output.

### GCI RAPID Kinetics for binding assay

The fragments were tested with Creoptix Wave (Malvern Panalyical) using PCH-NTA chips. First, the chip surface was conditioned with 1M NaCl, 0.1 M borate pH 9.0 buffer for 180 s, followed by an injection of 250 mM EDTA solution for 180 s. Then the chip surface was equilibrated by injecting running buffer (10 M HEPES pH 7.4, 150 mM NaCl, 0.05% Tween 20).

The immobilization of ZIKV NS2B-NS3 protease was performed using His capture combined with EDC/NHS conjugation. First, NiCl_2_ (500 µM) was injected to activate NTA groups on the chip surface for 420 s. Then, the running buffer was injected twice for 60 s to wash the chip surface. EDC/NHS (Xantec) was injected to activate carboxyl groups on the chip surface and to couple the target protease. The target protease was injected at a concentration of 15 µg/ml for 420 s on the FC2, FC3, and FC4 channels. Excessively activated groups were blocked with the injection of ethanolamine (1M pH 8.5, Xantec) solution. The FC1 blank surface was treated the same as the FC2/FC3/FC4 active channel except for the NS2B-NS3 protein capture step. The final NS2B-NS3 capture level was about 9000 pg/mm2 on the active channels. All steps within the immobilization were performed at 10 µL/min flow rate. Kinetic binding assay was performed through Repeated Analyte Pulses of Increasing Duration (RAPID) kinetics assay in Creoptix. Samples were injected at 250 µM for 5 s association and 20 s dissociation at 400 µL/min flow rate. Positive control compound was tested at 1µM for accurate result. DMSO correction was included in the method and injected as 0.5% DMSO addition to the running buffer +2%DMSO. Blank solution (the running buffer +2%DMSO) was injected after every 5th sample. All steps were conducted at 25°C. Data analysis was carried out with adjustment of X and y offset, DMSO calibration, and blank subtraction in WAVE control software 4. 5.18.

### Fluorescence-dose-response inhibitory activity assay

This protocol was adapted from a previous study^25^ with minor modifications. Compounds were seeded into assay-ready plates (Greiner 384 low volume, cat. no. 784076) using an Echo 555 acoustic dispenser, and dimethyl sulfoxide (DMSO) was back-filled for a uniform concentration in assay plates (DMSO concentration 1.5%). Dose-response assays were performed in 12-point dilutions of twofold, starting at 300 μM. Reagents for the assay were dispensed into the assay plate in 10 μl volumes for a final volume of 20 μl.

Final reaction concentrations were 20 mM Tris pH 8.5, 0.01% Triton, 10% glycerol, 15 nM ZIKV NS2B-NS3 and 5 µM fluorogenic peptide substrate (Boc-Gly-Arg-Arg-AMC, CAS # [113866-14-1(free base)], Biosynth (FB110553)). ZIKV NS2B-NS3 protease was pre-incubated for 2 hr followed by the addition of substrate and a further 30 mins incubation with the substrate (all incubations performed at room temperature).

Protease reaction was measured in a BMG Pherastar FS with a 360/470 nm excitation/emission filter set. Raw data were mapped and normalized to high (Protease with DMSO, no compounds) and low (No Protease, no compounds) controls using Genedata Screener software. Normalized data were then uploaded to CDD Vault (Collaborative Drug Discovery). Dose-response curves were generated for IC_50_ using nonlinear regression with the Levenberg–Marquardt algorithm with minimum inhibition 0% and maximum inhibition 100%. To each run we added the reported peptide-hybrid inhibitor compound **36** as a positive control (for a more accurate IC_50_ measurements of this compound, dose response started from 100 µM).

## Data availability

Crystallographic coordinate and structure factor for cZiPro crystal structure has been deposited in the PDB with the accession code 8PN6. PDB codes of crystallographic fragment binders are listed in Crystallographic Supplementary Table.

## Acknowledgement

Research reported in this publication was supported by NIAID of the National Institute of Health under award number U19AI171399. The content is solely the responsibility of the authors and does not necessarily represent the official views of the National Institute of Health. Authors acknowledge Diamond Light Source for access to the I04-1 beamline and XChem facilities through proposal lb32627.

## Author contributions

X.N.: conceptualization, X-Ray data curation, formal analysis, visualization, writing—original draft, writing—review and editing; R.B.R.: All DMS data curation and analysis, review and editing; A.S.G: crystallography support, writing—review and editing; M.P.F.: Fragment progressing; C.K., J.S., and W.W.H.: DMS analysis, visualization; B.H.B., P.G.M., C.W.E.T.: XChem support. M.F., S.W., E.P.W., C.G.: Protein engineering, protein production and quality control; N.L., E.C., H.B.: biochemical and biophysical assays. R.M.L., M.W., W.T., A.V.C., M.W.: software, manuscript reviewing. J.C.A., D.F., L.K., K.K.: manuscript review and editing; M.J.E.: conceptualization, writing—review and editing; F.v.D.: funding acquisition, conceptualization, writing—review and editing.

## Reference

(1) de Araujo, T. V. B.; Ximenes, R. A. A.; Miranda-Filho, D. B.; Souza, W. V.; Montarroyos, U. R.; de Melo, A. P. L.; Valongueiro, S.; de Albuquerque, M.; Braga, C.; Filho, S. P. B.;, et al. Association between microcephaly, Zika virus infection, and other risk factors in Brazil: final report of a case-control study. Lancet Infect Dis 2018, 18 (3), 328–336. DOI: 10.1016/S1473-3099(17)30727-2 From NLM Medline.

(2) Krauer, F.; Riesen, M.; Reveiz, L.; Oladapo, O. T.; Martinez-Vega, R.; Porgo, T. V.; Haefliger, A.; Broutet, N. J.; Low, N.; Group, W. H. O. Z. C. W. Zika Virus Infection as a Cause of Congenital Brain Abnormalities and Guillain-Barre Syndrome: Systematic Review. PLoS Med 2017, 14 (1), e1002203. DOI: 10.1371/journal.pmed.1002203 From NLM Medline.

(3) Lowe, R.; Barcellos, C.; Brasil, P.; Cruz, O. G.; Honorio, N. A.; Kuper, H.; Carvalho, M. S. The Zika Virus Epidemic in Brazil: From Discovery to Future Implications. Int J Environ Res Public Health 2018, 15 (1). DOI: 10.3390/ijerph15010096 From NLM Medline.

(4) Saiz, J. C. Therapeutic Advances Against ZIKV: A Quick Response, a Long Way to Go. Pharmaceuticals (Basel) 2019, 12 (3). DOI: 10.3390/ph12030127 From NLM PubMed-not-MEDLINE.

(5) Chappell, K. J.; Stoermer, M. J.; Fairlie, D. P.; Young, P. R. Insights to substrate binding and processing by West Nile Virus NS3 protease through combined modeling, protease mutagenesis, and kinetic studies. J Biol Chem 2006, 281 (50), 38448–38458. DOI: 10.1074/jbc.M607641200 From NLM Medline.

(6) Zhang, Z.; Li, Y.; Loh, Y. R.; Phoo, W. W.; Hung, A. W.; Kang, C.; Luo, D. Crystal structure of unlinked NS2B-NS3 protease from Zika virus. Science 2016, 354 (6319), 1597–1600. DOI: 10.1126/science.aai9309 From NLM Medline.

(7) Chen, X.; Yang, K.; Wu, C.; Chen, C.; Hu, C.; Buzovetsky, O.; Wang, Z.; Ji, X.; Xiong, Y.; Yang, H. Mechanisms of activation and inhibition of Zika virus NS2B-NS3 protease. Cell Res 2016, 26 (11), 1260–1263. DOI: 10.1038/cr.2016.116 From NLM Medline.

(8) Erbel, P.; Schiering, N.; D’Arcy, A.; Renatus, M.; Kroemer, M.; Lim, S. P.; Yin, Z.; Keller, T. H.; Vasudevan, S. G.; Hommel, U. Structural basis for the activation of flaviviral NS3 proteases from dengue and West Nile virus. Nat Struct Mol Biol 2006, 13 (4), 372–373. DOI: 10.1038/nsmb1073 From NLM Medline.

(9) Meewan, I.; Shiryaev, S. A.; Kattoula, J.; Huang, C. T.; Lin, V.; Chuang, C. H.; Terskikh, A. V.; Abagyan, R. Allosteric Inhibitors of Zika Virus NS2B-NS3 Protease Targeting Protease in “Super-Open” Conformation. Viruses 2023, 15 (5). DOI: 10.3390/v15051106 From NLM Medline.

(10) Braun, N. J.; Quek, J. P.; Huber, S.; Kouretova, J.; Rogge, D.; Lang-Henkel, H.; Cheong, E. Z. K.; Chew, B. L. A.; Heine, A.; Luo, D.; Steinmetzer, T. Structure-Based Macrocyclization of Substrate Analogue NS2B-NS3 Protease Inhibitors of Zika, West Nile and Dengue viruses. ChemMedChem 2020, 15 (15), 1439–1452. DOI: 10.1002/cmdc.202000237 From NLM Medline.

(11) Huber, S.; Braun, N. J.; Schmacke, L. C.; Quek, J. P.; Murra, R.; Bender, D.; Hildt, E.; Luo, D.; Heine, A.; Steinmetzer, T. Structure-Based Optimization and Characterization of Macrocyclic Zika Virus NS2B-NS3 Protease Inhibitors. J Med Chem 2022, 65 (9), 6555–6572. DOI: 10.1021/acs.jmedchem.1c01860 From NLM Medline.

(12) Nunes, D. A. F.; Santos, F.; da Fonseca, S. T. D.; de Lima, W. G.; Nizer, W.; Ferreira, J. M. S.; de Magalhaes, J. C. NS2B-NS3 protease inhibitors as promising compounds in the development of antivirals against Zika virus: A systematic review. J Med Virol 2022, 94 (2), 442–453. DOI: 10.1002/jmv.27386 From NLM Medline.

(13) Yildiz, M.; Ghosh, S.; Bell, J. A.; Sherman, W.; Hardy, J. A. Allosteric inhibition of the NS2B-NS3 protease from dengue virus. ACS Chem Biol 2013, 8 (12), 2744–2752. DOI: 10.1021/cb400612h From NLM Medline.

(14) Li, Z.; Brecher, M.; Deng, Y. Q.; Zhang, J.; Sakamuru, S.; Liu, B.; Huang, R.; Koetzner, C. A.; Allen, C. A.; Jones, S. A.;, et al. Existing drugs as broad-spectrum and potent inhibitors for Zika virus by targeting NS2B-NS3 interaction. Cell Res 2017, 27 (8), 1046–1064. DOI: 10.1038/cr.2017.88 From NLM Medline.

(15) Brecher, M.; Li, Z.; Liu, B.; Zhang, J.; Koetzner, C. A.; Alifarag, A.; Jones, S. A.; Lin, Q.; Kramer, L. D.; Li, H. A conformational switch high-throughput screening assay and allosteric inhibition of the flavivirus NS2B-NS3 protease. PLoS Pathog 2017, 13 (5), e1006411. DOI: 10.1371/journal.ppat.1006411 From NLM Medline.

(16) Burton, T. D.; Eyre, N. S. Applications of Deep Mutational Scanning in Virology. Viruses 2021, 13 (6). DOI: 10.3390/v13061020 From NLM Medline.

(17) Dadonaite, B.; Brown, J.; McMahon, T. E.; Farrell, A. G.; Figgins, M. D.; Asarnow, D.; Stewart, C.; Lee, J.; Logue, J.; Bedford, T.;, et al. Spike deep mutational scanning helps predict success of SARS-CoV-2 clades. Nature 2024, 631 (8021), 617–626. DOI: 10.1038/s41586-024-07636-1 From NLM Medline.

(18) Sourisseau, M.; Lawrence, D. J. P.; Schwarz, M. C.; Storrs, C. H.; Veit, E. C.; Bloom, J. D.; Evans, M. J. Deep Mutational Scanning Comprehensively Maps How Zika Envelope Protein Mutations Affect Viral Growth and Antibody Escape. J Virol 2019, 93 (23). DOI: 10.1128/JVI.01291-19 From NLM Medline.

(19) Phoo, W. W.; Li, Y.; Zhang, Z. Z.; Lee, M. Y. Q.; Loh, Y. R.; Tan, Y. B.; Ng, E. Y.; Lescar, J.; Kang, C. B.; Luo, D. H. Structure of the NS2B-NS3 protease from Zika virus after self-cleavage. Nature Communications 2016, 7. DOI: ARTN 13410 10.1038/ncomms13410.

(20) Schwarz, M. C.; Sourisseau, M.; Espino, M. M.; Gray, E. S.; Chambers, M. T.; Tortorella, D.; Evans, M. J. Rescue of the 1947 Zika Virus Prototype Strain with a Cytomegalovirus Promoter-Driven cDNA Clone. mSphere 2016, 1(5). DOI: 10.1128/mSphere.00246-16 From NLM PubMed-not-MEDLINE.

(21) Blight, K. J.; Kolykhalov, A. A.; Rice, C. M. Efficient initiation of HCV RNA replication in cell culture. Science 2000, 290 (5498), 1972–1974. DOI: 10.1126/science.290.5498.1972 From NLM Medline.

(22) Kirkpatrick, E.; Qiu, X.; Wilson, P. C.; Bahl, J.; Krammer, F. The influenza virus hemagglutinin head evolves faster than the stalk domain. Sci Rep 2018, 8 (1), 10432. DOI: 10.1038/s41598-018-28706-1 From NLM Medline.

(23) Shiryaev, S. A.; Farhy, C.; Pinto, A.; Huang, C. T.; Simonetti, N.; Elong Ngono, A.; Dewing, A.; Shresta, S.; Pinkerton, A. B.; Cieplak, P.;, et al. Characterization of the Zika virus two-component NS2B-NS3 protease and structure-assisted identification of allosteric small-molecule antagonists. Antiviral Res 2017, 143, 218–229. DOI: 10.1016/j.antiviral.2017.04.015 From NLM Medline.

(24) Yao, Y.; Huo, T.; Lin, Y. L.; Nie, S.; Wu, F.; Hua, Y.; Wu, J.; Kneubehl, A. R.; Vogt, M. B.; Rico-Hesse, R.; Song, Y. Discovery, X-ray Crystallography and Antiviral Activity of Allosteric Inhibitors of Flavivirus NS2B-NS3 Protease. J Am Chem Soc 2019, 141 (17), 6832–6836. DOI: 10.1021/jacs.9b02505 From NLM Medline.

(25) Boby, M. L.; Fearon, D.; Ferla, M.; Filep, M.; Koekemoer, L.; Robinson, M. C.; dagger, C. M. C.; Chodera, J. D.; Lee, A. A.; London, N.;, et al. Open science discovery of potent noncovalent SARS-CoV-2 main protease inhibitors. Science 2023, 382 (6671), eabo7201. DOI: 10.1126/science.abo7201 From NLM Medline.

(26) Starvaggi, J.; Previti, S.; Zappala, M.; Ettari, R. The Inhibition of NS2B/NS3 Protease: A New Therapeutic Opportunity to Treat Dengue and Zika Virus Infection. Int J Mol Sci 2024, 25 (8). DOI: 10.3390/ijms25084376 From NLM Medline.

(27) Schuller, M.; Correy, G. J.; Gahbauer, S.; Fearon, D.; Wu, T.; Diaz, R. E.; Young, I. D.; Carvalho Martins, L.; Smith, D. H.; Schulze-Gahmen, U.;, et al. Fragment binding to the Nsp3 macrodomain of SARS-CoV-2 identified through crystallographic screening and computational docking. Sci Adv 2021, 7(16). DOI: 10.1126/sciadv.abf8711 From NLM Medline.

(28) Schiebel, J.; Radeva, N.; Koster, H.; Metz, A.; Krotzky, T.; Kuhnert, M.; Diederich, W. E.; Heine, A.; Neumann, L.; Atmanene, C.;, et al. One Question, Multiple Answers: Biochemical and Biophysical Screening Methods Retrieve Deviating Fragment Hit Lists. ChemMedChem 2015, 10 (9), 1511–1521. DOI: 10.1002/cmdc.201500267 From NLM Medline.

(29) Ferla MP, S.-G. R., Skyner RE, Gahbauer S, Taylor JC, von Delft F, et al. Fragmenstein: predicting protein-ligand structures of compounds derived from known crystallographic fragment hits using a strict conserved-binding–based methodology. ChemRxiv. 2024. DOI: 10.26434/chemrxiv-2024-17w01.

(30) Ali, A.; Bandaranayake, R. M.; Cai, Y.; King, N. M.; Kolli, M.; Mittal, S.; Murzycki, J. F.; Nalam, M. N. L.; Nalivaika, E. A.; Ozen, A.;, et al. Molecular Basis for Drug Resistance in HIV-1 Protease. Viruses 2010, 2 (11), 2509–2535. DOI: 10.3390/v2112509 From NLM PubMed-not-MEDLINE.

(31) Santos, N. P.; Santos, L. H.; de Magalhaes, M. T. Q.; Lei, J.; Hilgenfeld, R.; Ferreira, R. S.; Bleicher, L. Characterization of an Allosteric Pocket in Zika Virus NS2B-NS3 Protease. J Chem Inf Model 2022, 62 (4), 945–957. DOI: 10.1021/acs.jcim.1c01326.

(32) Fearon, D., Powell, A., Douangamath, A., Dias, A., Tomlinson, C.W.E., Balcomb, B.H., Aschenbrenner, J.C., Aimon, A., Barker, I.A., Brandão-Neto, J., Collins, P., Dunnett, L.E., Fairhead, M., Gildea, R.J., Golding, M., Gorrie-Stone, T., Hathaway, P.V., Koekemoer, L., Krojer, T., Lithgo, R.M., Maclean, E.M., Marples, P.G., Ni, X., Skyner, R., Talon, R., Thompson, W., Wild, C.F., Winokan, M., Wright, N.D., Winter, G., Shotton, E.J. and von Delft, F. . Accelerating Drug Discovery With High-Throughput Crystallographic Fragment Screening and Structural Enablement. Applied Research 2025, 4. DOI: 10.1002/appl.202400192.

(33) (33) Gahbauer, S.; Correy, G. J.; Schuller, M.; Ferla, M. P.; Doruk, Y. U.; Rachman, M.; Wu, T.; Diolaiti, M.; Wang, S.; Neitz, R. J.; et al. Iterative computational design and crystallographic screening identifies potent inhibitors targeting the Nsp3 Macrodomain of SARS-CoV-2. bioRxiv 2022. DOI: 10.1101/2022.06.27.497816 From NLM PubMed-not-MEDLINE.

(34) Michael Fairhead; Lizbe Koekemoer; Eleanor Williams; Frank von Delft. A selection of Golden Gate vectors to simplify recombinant protein production in Escherichia coli. BioRxiv 2024, PrePrint. DOI: DOI: 10.1101/2024.02.13.579886.

(35) Cox, O. B.; Krojer, T.; Collins, P.; Monteiro, O.; Talon, R.; Bradley, A.; Fedorov, O.; Amin, J.; Marsden, B. D.; Spencer, J.;, et al. A poised fragment library enables rapid synthetic expansion yielding the first reported inhibitors of PHIP(2), an atypical bromodomain. Chem Sci 2016, 7 (3), 2322–2330. DOI: 10.1039/c5sc03115j.

(36) O’Reilly, M.; Cleasby, A.; Davies, T. G.; Hall, R. J.; Ludlow, R. F.; Murray, C. W.; Tisi, D.; Jhoti, H. Crystallographic screening using ultra-low-molecular-weight ligands to guide drug design. Drug Discov Today 2019, 24 (5), 1081–1086. DOI: 10.1016/j.drudis.2019.03.009.

(37) Keeley, A.; Abranyi-Balogh, P.; Keseru, G. M. Design and characterization of a heterocyclic electrophilic fragment library for the discovery of cysteine-targeted covalent inhibitors. Medchemcomm 2019, 10 (2), 263–267. DOI: 10.1039/c8md00327k From NLM PubMed-not-MEDLINE.

(38) Bajusz, D.; Wade, W. S.; Satala, G.; Bojarski, A. J.; Ilas, J.; Ebner, J.; Grebien, F.; Papp, H.; Jakab, F.; Douangamath, A.;, et al. Exploring protein hotspots by optimized fragment pharmacophores. Nature Communications 2021, 12 (1). DOI: ARTN 3201 10.1038/s41467-021-23443-y.

(39) Wright, N. D.; Collins, P.; Koekemoer, L.; Krojer, T.; Talon, R.; Nelson, E.; Ye, M.; Nowak, R.; Newman, J.; Ng, J. T.;, et al. The low-cost Shifter microscope stage transforms the speed and robustness of protein crystal harvesting. Acta Crystallogr D Struct Biol 2021, 77 (Pt 1), 62–74. DOI: 10.1107/S2059798320014114 From NLM Medline.

(40) Kabsch, W. Xds. Acta Crystallogr D Biol Crystallogr 2010, 66 (Pt 2), 125–132. DOI: 10.1107/S0907444909047337 From NLM Medline.

(41) Vonrhein, C.; Flensburg, C.; Keller, P.; Sharff, A.; Smart, O.; Paciorek, W.; Womack, T.; Bricogne, G. Data processing and analysis with the autoPROC toolbox. Acta Crystallogr D Biol Crystallogr 2011, 67 (Pt 4), 293–302. DOI: 10.1107/S0907444911007773 From NLM Medline.

(42) Winter, G.: an expert system for macromolecular crystallography data reduction. J Appl Crystallogr 2010, 43, 186–190. DOI: 10.1107/S0021889809045701.

(43) Winter, G.; Waterman, D. G.; Parkhurst, J. M.; Brewster, A. S.; Gildea, R. J.; Gerstel, M.; Fuentes-Montero, L.; Vollmar, M.; Michels-Clark, T.; Young, I. D.;, et al. DIALS: implementation and evaluation of a new integration package. Acta Crystallogr D Struct Biol 2018, 74 (Pt 2), 85–97. DOI: 10.1107/S2059798317017235 From NLM Medline.

(44) Krojer, T.; Talon, R.; Pearce, N.; Collins, P.; Douangamath, A.; Brandao-Neto, J.; Dias, A.; Marsden, B.; von Delft, F. The XChemExplorer graphical workflow tool for routine or large-scale protein-ligand structure determination. Acta Crystallogr D Struct Biol 2017, 73 (Pt 3), 267–278. DOI: 10.1107/S2059798316020234 From NLM Medline.

(45) Long, F.; Nicholls, R. A.; Emsley, P.; Graaeulis, S.; Merkys, A.; Vaitkus, A.; Murshudov, G. N. AceDRG: a stereochemical description generator for ligands. Acta Crystallogr D Struct Biol 2017, 73 (Pt 2), 112–122. DOI: 10.1107/S2059798317000067 From NLM Medline.

(46) Smart, O. S., Womack, T. O., Sharff, A., Flensburg, C., Keller, P., Paciorek, W., Vonrhein, C. and Bricogne, G. (2011). Grade, version 1.2.20. Cambridge, United Kingdom, Global Phasing Ltd., . (accessed.

(47) Pearce, N. M.; Krojer, T.; Bradley, A. R.; Collins, P.; Nowak, R. P.; Talon, R.; Marsden, B. D.; Kelm, S.; Shi, J.; Deane, C. M.; von Delft, F. A multi-crystal method for extracting obscured crystallographic states from conventionally uninterpretable electron density. Nat Commun 2017, 8, 15123. DOI: 10.1038/ncomms15123 From NLM PubMed-not-MEDLINE.

(48) Keegan, R.; Wojdyr, M.; Winter, G.; Ashton, A. DIMPLE: A difference map pipeline for the rapid screening of crystals on the beamline. Acta Crystallogr A 2015, 71, S18–S18. DOI: 10.1107/S2053273315099702.

(49) Emsley, P.; Lohkamp, B.; Scott, W. G.; Cowtan, K. Features and development of Coot. Acta Crystallogr D Biol Crystallogr 2010, 66 (Pt 4), 486–501. DOI: 10.1107/S0907444910007493 From NLM Medline.

(50) Smart, O. S.; Womack, T. O.; Flensburg, C.; Keller, P.; Paciorek, W.; Sharff, A.; Vonrhein, C.; Bricogne, G. Exploiting structure similarity in refinement: automated NCS and target-structure restraints in BUSTER. Acta Crystallogr D Biol Crystallogr 2012, 68 (Pt 4), 368–380. DOI: 10.1107/S0907444911056058 From NLM Medline.

(51) Crill, W. D.; Chang, G. J. Localization and characterization of flavivirus envelope glycoprotein cross-reactive epitopes. J Virol 2004, 78 (24), 13975–13986. DOI: 10.1128/JVI.78.24.13975-13986.2004 From NLM Medline.

(52) Bloom, J. D. An experimentally determined evolutionary model dramatically improves phylogenetic fit. Mol Biol Evol 2014, 31 (8), 1956–1978. DOI: 10.1093/molbev/msu173 From NLM Medline.

(53) Fulton, B. O.; Sachs, D.; Schwarz, M. C.; Palese, P.; Evans, M. J. Transposon Mutagenesis of the Zika Virus Genome Highlights Regions Essential for RNA Replication and Restricted for Immune Evasion. J Virol 2017, 91 (15). DOI: 10.1128/JVI.00698-17 From NLM Medline.

(54) Doud, M. B.; Bloom, J. D. Accurate Measurement of the Effects of All Amino-Acid Mutations on Influenza Hemagglutinin. Viruses 2016, 8 (6). DOI: 10.3390/v8060155 From NLM Medline.

(55) Bloom, J. D. Software for the analysis and visualization of deep mutational scanning data. BMC Bioinformatics 2015, 16, 168. DOI: 10.1186/s12859-015-0590-4 From NLM Medline.

