## Supplementary material for "Crystallographic fragment screening and deep mutational scanning of Zika virus NS2B-NS3 protease enable development of resistance-resilient inhibitors": Supplementary information.docx

**Supplementary Table 1.** Data collection and refinement statistics

| Protein | ZIKV NS2B-NS3 |
| --- | --- |
| PDB accession code | 8PN6 |
| Data Collection |  |
| Resolution^a^ (Å) | 33.70 – 1.60 (1.63 -1.60) |
| Spacegroup | *P* 4_3_22 |
| Cell dimensions | *a* = 43, b = 43, *c* = 216.6 Å |
|  | *α= β* = *γ* = 90.0° |
| No. unique reflections^a^ | 28203 (1313) |
| Completeness^a^ (%) | 100.0 (99.9) |
| I/σI^a^ | 24.6 (3.4) |
| Rpim^a^ | 0.016 (0.310) |
| CC (1/2) | 0.999 (0.977) |
| Redundancy^a^ | 22.4 (22.2) |
| Refinement |  |
| No. atoms in refinement | 1594 |
| Average B factor (Å^2^) | 41.0 |
| R_fact_ (%) | 20.8 |
| R_free_ (%) | 25.8 |
| rms deviation bond^b^ (Å) | 0.009 |
| rms deviation angle^b^ (°) | 1.37 |
| Molprobity Ramachandran Favored (%) | 96.86 |
| Outlier (%) | 0 |

^a^ Values in brackets show the statistics for the highest resolution shells.

^b^ rms indicates root-mean-square.

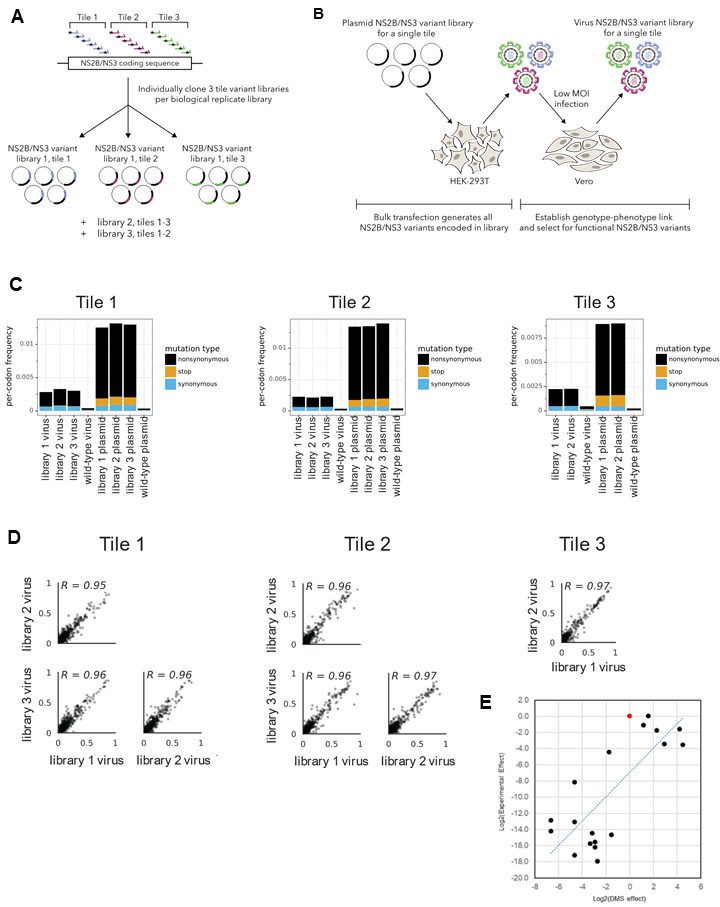

**Supplementary Figure1. Creation of mutant plasmid libraries and mutant virus of ZIKV NS2B-NS3 protease.** (A) Using a previously published workflow, we measured the tolerance and preference for all possible amino acid mutations to the entire coding sequence of ZIKV NS2B and NS3 protease over three tiled segments of the genome. Three biological replicates were created for each tile except tile 3 which had only two replicates. (B) HEK293T cells were transfected with variants from each tile to create a variant library. The resulting viruses were used to perform a low MOI infection on Vero cells to produce a phenotype-genotype link. (C) As expected, stop codons and non-synonymous mutations were purged from the library after passaging. (D) There is a high correlation between amino acid preferences measured with replicate deep mutational scanning libraries. (E) There is a good correlation between mutation effects measured by DMS and mutation effects measured experimentally (Pearson correlation coefficient of 0.769). Although some mutations were preferred over wildtype (red) in the DMS experiment, no mutations grew better than wildtype (red) at 72 hours post-transfection.
