## Supplementary material for "Crystallographic fragment screening and deep mutational scanning of Zika virus NS2B-NS3 protease enable development of resistance-resilient inhibitors": Supplementary information_DMS experiment.docx

**Supplemental file for extended methods of deep mutational scanning**

Mutant plasmid libraries were cloned into our previously described single-plasmid reverse genetics system for ZIKV strain MR766^1^ (sequence is available at Genbank accession KX830961) using techniques like those described in our previous ZIKV envelope protein DMS paper^2^. In this plasmid, unique XhoI and BstBI restriction sites flank the NS2B-NS3 protease coding sequence. In order to enable efficient cloning into this plasmid backbone, we designed a recipient plasmid in which this 1,119 nucleotide region was replaced by an inert stuffer fragment (sequence available upon request). Any contamination of this vector in our mutant libraries would not result in infectious virus production from the vector.

We created NS2B-NS3 protease codon-mutant libraries using a previously described PCR mutagenesis approach^3^. First, we designed sets of forward and reverse oligos that randomized each codon with an NNK sequence where N is any nucleotide and K is either a G or a T. These primers were designed using a Python script (available at https://github.com/jbloomlab/CodonTilingPrimers) to have an average melting temperature of about ~65°C^4^. We designed 308 forward and reverse oligo pairs targeting the entire 130 amino acids of NS2B and the 178 amino acids of NS3 comprising its protease domain. To ensure that we could maintain library complexity at all stages of the screen, we subdivided our NS2B-NS3 coding sequencing of interest into three separate tiling DMS library segments, hereinafter referred to simply as “tiles.” The first and second tiles each cover 103 mutant codons, and the third tile covers 102 mutant codons. One added benefit to this tile size was that it fit well into a single paired-end Illumina read, which simplified the deep sequencing and analysis. For each tile, we created three individual libraries to be screened as independent biological replicates. Forward and reverse oligos for each tile were synthesized as oPools Oligo Pools (Integrated DNA Technologies).

We performed the mutagenesis PCR using these primers and the following end primers flanking the NS2B/NS3 protease coding sequence that provide Gibson Assembly overlaps with the XhoI and BstBI restriction sites: 5’-TACCAATCTTGGCTGCTCTAACACCACTAGCTC -3’ and 5’-AGTTAGCTGCTTCTTCTTCAGCATCGAGGGTTC -3’. Two rounds of mutagenic PCR were performed in triplicate using a similar approach to previous work^3^ with the following adjustments: For the reaction with the forward oPool oligos, 12 ng of WT NS2B/NS3 PCR product was used as template, with 2 µL of the reverse primer at 5 µM, and 2 µL of the forward oPool mixture at 1 µM; For the reaction with the reverse oPool oligos, 12ng of PCR product was used as template, with 2 µL of 5 µM forward primer, and 1 µL of the “ANN” and “CNN” at 2 µM was used. Due to technical restraints, the reverse NNK pools must be synthesized as two separate libraries, one representing all “ANN” mutants and one representing all “CNN” mutants. These were used in a standard PCR reaction using Q5 High-Fidelity DNA Polymerase (New England Biolabs, Ipswich, MA) in a final volume of 30 µL. The PCR template was amplified using the program of (1) 98°C for 2:00; (2) 98°C for 0:10; (3) 70°C for 0:01; (4) cool at 0.5°C/sec to 50°C and hold for 0:30; (5) 70°C for 1:30; 6) Go to (2) (3 total cycles); (7) 70°C 2:00; (8) 4°C hold. Then, the forward and reverse PCR reactions were diluted 1:4, and 4 µL of these dilutions were used in a second round of PCR, with 2 µL of the forward and reverse primers above at 5 µM in a 30 µL PCR reaction. The PCR program was identical to that above except it was used for 20 total cycles. The resulting PCR products were gel purified using an E.Z.N.A. Gel Extraction Kit (Omega Biotek, Norcross, GA). After XhoI and BstBI digestion and purification of the recipient vector plasmid, mutagenized NS2B-NS3 protease amplicons were cloned into the recipient MR766 genome using Gibson assembly and transformed into 10-beta electrocompetent E. coli (NEB). Transformants were plated on LB supplemented with ampicillin for counting colony numbers and in bottle culture of LB supplemented with carbenicillin for full library growth^5, 6^. To help ensure higher plasmid stability, the bacterial growth steps were performed at 30°C. Each codon mutant library was prepared by HiSpeed Maxiprep (Qiagen) from transformants harvested from each bottle culture.

**Rescue and screening of ZIKV NS2B-NS3 protease mutant libraries.**

ZIKV NS2B/NS3 protease DMS libraries were produced by transfecting plasmids into HEK 293T cells using a protocol to maintain library complexity as previously reported^2^. Wild type virus was rescued in parallel as a control. Briefly, prior to transfection, poly-lysine coated 6-well plates were seeded overnight with 4 × 105 293T cells. For each library replicate, all wells of a 6-well plate were transfected with 2 μg of plasmid DNA per well and the TransIT-LT1 transfection reagent (Mirus Bio, Madison WI) following the manufacturer's recommendations. The viral supernatants were collected at day 2 and 3 post transfection, pooled, centrifuged to remove cellular debris, and stored at −80°C.

**Titration of infectious viruses and ZIKV NS2B-NS3 libraries.**

The infectious titers of rescued wild type virus and mutant virus libraries were determined by immunostaining with the pan-flavivirus E protein reactive 4G2 antibody and flow cytometry, as previously described^7^. Briefly, one day prior to infection, 24-well plates were seeded at a density of 1 x105 cells/well. The next day, 250 uL of serially diluted transfection supernatants in DMEM with 3% FBS was added to each well (in duplicate). At 24 hours post infection, at which time initial infection events would be detectable but secondary infections would not be, cells were fixed in 4% paraformaldehyde and stained with 4G2 antibodies and goat-anti-human secondary antibody conjugated to Alexa Fluor 647 (Thermo Fisher Scientific, Waltham, MA). Data were acquired on an Attune flow cytometer (Thermo Fisher Scientific, Waltham, MA) and analyzed using FlowJo (Tree Star, USA). Only the viral dilutions leading to less than 20% of infected cells were used to calculate the infectious titers. Infectious units per mL were calculated using the following formula: “percentage of infectious events x number of cells / volume (mL) of viral inoculum”.

**Deep sequencing using barcoded-subamplicon sequencing**

Total RNA for each tile within the NS2B/NS3 protease coding sequence was reverse transcribed using AccuScript High Fidelity Reverse Transcriptase (Agilent Technologies, La Jolla, CA) with primers for tile 1 (5’ - TTACTCACAAGGAGTGGGAAG - 3’ and 5’ - CATGCCACAGATGGCCATCAG - 3’), tile 2 (5’ – GAGATCATACTCAAGGTGGTC - 3’ and 5’ - AGGCCCACAGTATGACACCAA - 3’), and tile 3 (5’ – TGGGGGGATGTCAAGCAGGAC - 3’ and 5’ - CAGCATCGAGGGTTCGAAACA - 3’). Following the manufacturer’s instructions, we used 1 ug of RNA for each sample. To obtain high sequencing accuracy, we followed the barcoded-subamplicon sequencing approach described^8^ (see also https://jbloomlab.github.io/dms_tools2/bcsubamp.html). We generated amplicons for each of the NS2B/NS3 protease tiles using the same RT primers listed above, followed by two barcoding PCRs. For the amplicon PCR, we used 2 µL of the RT reaction or 10ng for amplification of the plasmid libraries. The cycling program used for the amplicon PCR was: (1) 98°C for 2:00; (2) 98°C for 0:10; (3) 70°C for 0:01; (4) cool at 0.5°C/sec to 50°C and hold for 0:30; (5) 70°C for 0:30; (6) Go to (2) (20 total cycles); (7) 70°C 2:00; (8) 4°C hold. The first primer used for amplification adds an N8 randomized barcode and the second primer adds Illumina adapter sequences. Round 1 PCR reactions were set up using Q5 Polymerase, 4 ng of amplicon PCR product, 2 μL of forward primer diluted to 5 μM, and 2 μL of reverse primer diluted to 5 μM. The primers used in Round 1 PCR for tile 1 (5’ - CTTTCCCTACACGACGCTCTTCCGATCTNNNNNNNNTTACTCACAAGGAGTGGGAAG - 3’ and 5’ - GGAGTTCAGACGTGTGCTCTTCCGATCTNNNNNNNNCATGCCACAGATGGCCATCAG - 3’), tile 2 (5’ - CTTTCCCTACACGACGCTCTTCCGATCTNNNNNNNNGAGATCATACTCAAGGTGGTC -3’ and 5’ - GGAGTTCAGACGTGTGCTCTTCCGATCTNNNNNNNNAGGCCCACAGTATGACACCAA - 3’), and for tile 3 (5’ - CTTTCCCTACACGACGCTCTTCCGATCTNNNNNNNNTGGGGGGATGTCAAGCAGGAC - 3’ and 5’ - GGAGTTCAGACGTGTGCTCTTCCGATCTNNNNNNNNCAGCATCGAGGGTTCGAAACA - 3’) are listed. The cycling program used for the Round 1 PCR was: (1) 98°C for 2:00; (2) 98°C for 0:10; (3) 70°C for 0:01; (4) cool at 0.5°C/sec to 50°C and hold for 0:30; (5) 70°C for 0:30; (6) Go to (2) (9 total cycles); (7) 70°C 2:00; (8) 95°C for 1:00; (9) 4°C hold. The final denaturation step ensures that double-stranded DNA molecules entering Round 2 PCR will contain two uniquely barcoded variants.

Amplicons were diluted and used as template for a second round of PCR using ~2×105 single-stranded template molecules per sample. The primers used for Round 2 PCR add sample-specific indices and the Illumina cluster-generating sequences. The PCRs used a universal forward primer (5’ - AATGATACGGCGACCACCGAGATCTACACTCTTTCCCTACACGACGCTCTTCCGATCT - 3’) and a sample-specific reverse primer that allows for indexing of the reads during sequencing with general format of (5’ - CAAGCAGAAGACGGCATACGAGATxxxxxxxxGTGACTGGAGTTCAGACGTGTGCTCTTCCGATCT -3’) where x represents an index-specific nucleotide. The Round 2 PCR reactions contained Q5 Polymerase, Round 1 DNA diluted to 2×105 ssDNA molecules per subamplicon, 4 μL of forward primer diluted to 5 μM, and 4 μL of reverse primer diluted to 5 μM. The cycling program used for Round 2 PCR was: (1) 98°C for 2:00; (2) 98°C for 0:10; (3) 70°C for 0:01; (4) cool at 0.5°C/sec to 50°C and hold for 0:30; (5) 70°C for 0:30; (6) Go to (2) (24 total cycles); (7) 70°C 2min; (8) 4°C Hold. After each PCR step (amplicon, Round 1 and Round 2), amplicons were purified using AMPure XP beads (Beckman Coulter, Brea, CA) (1x bead-to-sample volume for amplicon, and 1.6x bead-to-sample volume for Round 1 and Round 2). DNA concentration was quantified using Quant-iT PicoGreen dsDNA Assay Kit (Thermo Fisher Scientific, Waltham, MA). After barcoding, the Round 2 subamplicons were pooled, size selected by gel extraction, purified with an E.Z.N.A. Gel Extraction Kit (Omega Biotek, Norcross, GA), and purified with AMPure XP beads (1.6x bead-to-sample volume). Subamplicons were sequenced with an Illumina HiSeq2500 using 2 × 250 bp paired-end reads in Rapid Run mode. The raw deep sequencing data have been deposited in the Sequence Read Archive as BioProject PRJNA1125458.

**ZIKV Gluc validation of DMS results**

To validate the predicted fitness of mutants from the DMS screen, mutants were cloned into a pACYC177 plasmid. The expression plasmid pACYC177 MR766 contains a ZIKV MR766 genome with a Gaussia luciferase inserted between NS1 and NS2A under the control of a T7 promoter. These plasmids were linearized with ScaI and purified with a Zymo DCC-5 column. The linearized DNA was used as a template in a Promega T7 Express In vitro transcription reaction, with the addition of m7g cap. The reaction was purified with the EZNA RNA Kit and the products were aliquoted to 5µg per tube and stored in the -80C.

For each electroporation, Huh7.5 cells that were washed twice in ice cold PBS were resuspended at 1.5x10^7 cells/mL. Approximately 400µL (6x10^6 cells) were added to Eppendorf tubes and mixed with 5µg of RNA. The mixture was transferred to a 1mm gap cuvette and electroporated with the following conditions. Cells were rested for 10 minutes in the cuvettes, then transferred to 10mL of pre-warmed DMEM with 3% FBS.

Electroporated cells were seeded at a density of 80,000-100,000 cells per well in a 24-well plate, along with 100,000 non-electroporated cells to enhance spread of virus in the well. Cells were harvested 3 days post electroporation and luciferase activity was measured using a Renilla luciferase assay.

**Reference**

(1) Schwarz, M. C.; Sourisseau, M.; Espino, M. M.; Gray, E. S.; Chambers, M. T.; Tortorella, D.; Evans, M. J. Rescue of the 1947 Zika Virus Prototype Strain with a Cytomegalovirus Promoter-Driven cDNA Clone. *mSphere* **2016**, *1* (5). DOI: 10.1128/mSphere.00246-16 From NLM PubMed-not-MEDLINE.

(2) Sourisseau, M.; Lawrence, D. J. P.; Schwarz, M. C.; Storrs, C. H.; Veit, E. C.; Bloom, J. D.; Evans, M. J. Deep Mutational Scanning Comprehensively Maps How Zika Envelope Protein Mutations Affect Viral Growth and Antibody Escape. *J Virol* **2019**, *93* (23). DOI: 10.1128/JVI.01291-19 From NLM Medline.

(3) Bloom, J. D. An experimentally determined evolutionary model dramatically improves phylogenetic fit. *Mol Biol Evol* **2014**, *31* (8), 1956-1978. DOI: 10.1093/molbev/msu173 From NLM Medline.

(4) Dingens, A. S.; Haddox, H. K.; Overbaugh, J.; Bloom, J. D. Comprehensive Mapping of HIV-1 Escape from a Broadly Neutralizing Antibody. *Cell Host Microbe* **2017**, *21* (6), 777-787 e774. DOI: 10.1016/j.chom.2017.05.003 From NLM Medline.

(5) Kriegler, M. *Gene Transfer and Expression: A Laboratory Manual.*; 1990.

(6) Hanahan, D.; Jessee, J.; Bloom, F. R. Plasmid transformation of Escherichia coli and other bacteria. *Methods Enzymol* **1991**, *204*, 63-113. DOI: 10.1016/0076-6879(91)04006-a From NLM Medline.

(7) Crill, W. D.; Chang, G. J. Localization and characterization of flavivirus envelope glycoprotein cross-reactive epitopes. *J Virol* **2004**, *78* (24), 13975-13986. DOI: 10.1128/JVI.78.24.13975-13986.2004 From NLM Medline.

(8) Doud, M. B.; Bloom, J. D. Accurate Measurement of the Effects of All Amino-Acid Mutations on Influenza Hemagglutinin. *Viruses* **2016**, *8* (6). DOI: 10.3390/v8060155 From NLM Medline.
